# Functional connectome of brainstem nuclei involved in autonomic, limbic, pain and sensory processing in living humans from 7 Tesla resting state fMRI

**DOI:** 10.1101/2021.10.18.464861

**Authors:** Simone Cauzzo, Kavita Singh, Matthew Stauder, María Guadalupe García-Gomar, Nicola Vanello, Claudio Passino, Jeffrey Staab, Iole Indovina, Marta Bianciardi

**Affiliations:** Brainstem Imaging Laboratory, Department of Radiology, Athinoula A. Martinos Center for Biomedical Imaging, Massachusetts General Hospital and Harvard Medical School, Boston, MA, United States; Life Sciences Institute, Sant’Anna School of Advanced Studies, Pisa, Italy; Dipartimento di Ingegneria dell’Informazione, University of Pisa, Pisa, Italy; Fondazione Toscana Gabriele Monasterio, Pisa, Italy; Department of Psychiatry and Psychology, Mayo Clinic, Rochester, MN, United States; Department of Otorhinolaryngology – Head and Neck Surgery, Mayo Clinic, Rochester, MN, United States; Department of Biomedical and Dental Sciences and Morphofunctional Imaging, University of Messina, Italy; Laboratory of Neuromotor Physiology, IRCCS Santa Lucia Foundation, Rome, Italy; Division of Sleep Medicine, Harvard University, Boston, MA

**Keywords:** 7 Tesla, human functional connectome, brainstem, central autonomic network, vestibular network

## Abstract

Despite remarkable advances in mapping the functional connectivity of the cortex, the functional connectivity of subcortical regions is understudied in living humans. This is the case for brainstem nuclei that control vital processes, such as autonomic, limbic, nociceptive and sensory functions. This is because of the lack of precise brainstem nuclei localization, of adequate sensitivity and resolution in the deepest brain regions, as well as of optimized processing for the brainstem. To close the gap between the cortex and the brainstem, on 20 healthy subjects, we computed a correlation-based functional connectome of 15 brainstem nuclei involved in autonomic, limbic, nociceptive, and sensory function (superior and inferior colliculi, ventral tegmental area-parabrachial pigmented nucleus complex, microcellular tegmental nucleus-prabigeminal nucleus complex, lateral and medial parabrachial nuclei, vestibular and superior olivary complex, superior and inferior medullary reticular formation, viscerosensory motor nucleus, raphe magnus, pallidus, and obscurus, and parvicellular reticular nucleus – alpha part) with the rest of the brain. Specifically, we exploited 1.1mm isotropic resolution 7 Tesla resting-state fMRI, ad-hoc coregistration and physiological noise correction strategies, and a recently developed probabilistic template of brainstem nuclei. Further, we used 2.5mm isotropic resolution resting-state fMRI data acquired on a 3 Tesla scanner to assess the translatability of our results to conventional datasets. We report highly consistent correlation coefficients across subjects, confirming available literature on autonomic, limbic, nociceptive and sensory pathways, as well as high interconnectivity within the central autonomic network and the vestibular network. Interestingly, our results showed evidence of vestibulo-autonomic interactions in line with previous work. Comparison of 7 Tesla and 3 Tesla findings showed high translatability of results to conventional settings for brainstem-cortical connectivity and good yet weaker translatability for brainstem-brainstem connectivity. The brainstem functional connectome might bring new insight in the understanding of autonomic, limbic, nociceptive and sensory function in health and disease.

## Introduction

Vital processes such as autonomic, limbic, nociceptive and sensory functions are controlled by a group of brainstem nuclei and their brain networks, such as the central autonomic network (CAN) [Macefield and Henderson, 2019], the limbic network [Liddell et al., 2005], and pathways related to nociception [Napadow et al., 2019], as well as sensory processing, including vestibular pathways [Balaban, 2002; Balaban, 2004].

As opposed to the cortex and other subcortical areas, brainstem functional MRI (fMRI) networks have been understudied in living humans. Indeed, studies reporting the functional connectivity of brainstem nuclei in health and its pathognomonic changes in neurological disorders are sparse [Bär et al., 2016; Beliveau et al., 2015; Bianciardi et al., 2016; Del Cerro et al., 2020; Coulombe et al., 2016; Englot et al., 2018; Huang et al., 2019]. This is due to several reasons. First, the lack of a detailed probabilistic template of brainstem nuclei in living humans, which makes precise localization of brainstem nuclei difficult, and results less replicable [Fenske et al., 2020; Mäki-Marttunen and Espeseth, 2021], as opposed to cortical and subcortical studies, for which several parcellations are available [Desikan et al., 2006; Destrieux et al., 2010; Eickhoff et al., 2018]. Second, the need of high spatial resolution, high sensitivity and reasonably short repetition time (TR) to investigate the connectivity of these small nuclei with the whole brain, which are often unachievable on conventional MRI scanners. Third, adequate physiological noise reduction is needed to observe functional responses in these deep and tiny brain regions due to partial volumes with neighboring cerebrospinal fluid and vascular spaces [Beissner, 2015].

In this context, the need for a comprehensive description of the brainstem-brain connectivity in physiological conditions is evident. To this end, we mapped in living humans the functional connectome of 15 brainstem nuclei involved in autonomic, limbic, pain, and sensory functions, extracted from 7 Tesla resting-state functional MRI. Crucial to this goal were the availability of a novel probabilistic MRI template of 33 nuclei of the brainstem developed by our group [Bianciardi et al., 2016; Bianciardi et al., 2018; García-Gomar et al., 2019; Garcia Gomar et al., 2021; Singh et al., 2020b; Singh et al., 2020a], the use of ultra-high magnetic field MRI, which offered the necessary spatiotemporal resolution and sensitivity while maintaining whole-brain coverage, and of physiological noise correction methods. Further, we inspected the feasibility/limitations of mapping a similar brainstem connectome using conventional 3 Tesla data. Translatability to clinical studies is important to enable the investigation of pathological conditions involving autonomic, limbic, nociceptive and sensory functions, such as autonomic impairments [Henderson et al., 2006; Khazaie et al., 2017], psychiatric diseases [Köhler et al., 2019; Mochcovitch et al., 2014; Syan et al., 2018], and vestibular disorders [Roberts et al., 2018] .

## Methods

### Study participants and overview

The study protocol was approved by the Massachusetts General Hospital Institutional Review Board. After providing written informed consent in accordance with the declaration of Helsinki, 20 healthy subjects (10 males and 10 females; age 29.5±1.1 years) participated in two functional MRI sessions, one session at 7 Tesla (Magnetom, Siemens Healthineers, Erlangen, Germany), consisting of three resting-state runs (acquisition time per run = 10’07”), and another single-run session (acquisition time = 9’06”) at 3 Tesla (Connectom, Siemens Healthineers, Erlangen, Germany). The session order was randomized across subjects. Note that, for the purposes of this study, at 3 Tesla a conventional fMRI sequence was used, and the additional 3 Tesla Connectom scanner capabilities were not employed. A custom-built 64-channel receive coil and volume transmit coil was used at 3 Tesla [Keil et al., 2013]. During MRI acquisition, subjects were told to lay down and rest with their eyes closed. Foam pads placed between the subjects’ head and the MRI detector were used to minimize head motion during the acquisition.

### 7 Tesla Data Acquisition

A custom-built 32-channel receive coil and volume transmit coil was used at 7 Tesla. For each subject, three runs of 7 Tesla functional gradient-echo echo-planar images (EPIs) were acquired with the following parameters: isotropic voxel size = 1.1mm, matrix size = 180×240, GRAPPA factor = 3, nominal echo-spacing = 0.82ms, bandwidth = 1488Hz/Px, N. slices = 123, slice orientation = sagittal, slice-acquisition order = interleaved, echo time (TE) = 32ms, repetition time (TR) = 2.5s, flip angle (FA) = 75°, simultaneous-multi-slice factor = 3, N. repetitions = 210, phase-encoding direction = anterior-posterior, acquisition-time = 10’07”.

To perform geometric distortion correction of functional images, we acquired a fieldmap with the following parameters: isotropic voxel size = 2.0mm, matrix size = 116×132, bandwidth = 1515Hz/Px, N. slices = 80, slice orientation = sagittal, slice-acquisition order = interleaved, TE1 = 3.00ms, TE2 = 4.02ms, TR = 570.0ms, FA = 36°, simultaneous-multi-slice factor = 3, phase-encoding direction = anterior-posterior.

### 3 Tesla Data Acquisition

To assess the connectivity reproducibility using 3 Tesla MRI, on the same subjects, we acquired one run of conventional functional gradient-echo EPIs (isotropic voxel size = 2.5mm, matrix size = 100×215, GRAPPA factor = 2, nominal echo-spacing = 0.5ms, readout bandwidth = 2420Hz/Px, N. slices = 64, slice orientation = transversal, slice-acquisition order = interleaved, TE = 30ms, TR = 3.5s, FA = 85°, N. repetitions = 150, phase-encoding direction = anterior-posterior, acquisition time = 9’06”) and a fieldmap (isotropic voxel size = 2.5mm, FOV = 215×215, bandwidth = 300Hz/Px, N. slices = 128, slice orientation = sagittal, slice-acquisition order = interleaved, TE1 = 4.92ms, TE2 = 4.02ms, TR = 849.0ms, FA = 85°, simultaneous-multi-slice factor = 2’19”, phase-encoding direction = anterior-posterior). Note that, for the purposes of this study, at 3 Tesla a conventional fMRI sequence was used, and the additional 3 Tesla Connectom scanner capabilities were not employed. A custom-built 64-channel receive coil and volume transmit coil was used at 3 Tesla [Keil et al., 2013].

To parcellate the brain in cortical and subcortical regions, as well as to aid the coregistration of the fMRIs to stereotactic space, we also acquired an anatomical T1-weighted multi-echo MPRAGE (MEMPRAGE) image with parameters: isotropic voxel size = 1mm, TR = 2.53s, TEs = 1.69, 3.5, 5.3, 7.2ms, inversion-time = 1.5s, FA = 7°, FOV = 256×256×176mm^3^, bandwidth = 650Hz/Px, GRAPPA factor = 3, slice orientation = sagittal, slice-acquisition order = anterior-posterior, acquisition time = 4 28”.

### Preprocessing of MEMPRAGEs and fieldmaps

The root mean square across echo times was extracted from each MEMPRAGE image, the output was then rotated to standard orientation (‘RPI’). Bias-field correction was applied with SPM [Frackowiak et al., 1997] tools, then we used FSL routines to extract the brain and crop the image (FMRIB Software Library, FSL 5.0.7, Oxford, UK). Brain parcellations were generated on the MEMPRAGE with Freesurfer [Destrieux et al., 2010] to obtain cortical and subcortical targets. The preprocessed MEMPRAGEs were then iteratively aligned and averaged to build a group-based optimal template with the use of the Advanced Normalization Tool (ANTs, Philadelphia, PA, United States). The group-based optimal template was then registered to the MNI152_1mm template [Grabner et al., 2006] through an affine transformation and a nonlinear warp. Finally, we concatenated the transformation matrices from single subjects’ MEMPRAGEs to the optimal template and from the optimal template to MNI to obtain the full coregistration matrix, which aligns the MEMPRAGE to the MNI template.

Fieldmaps were mainly preprocessed in FSL, except for the SPM-based bias field correction: specifically, the magnitude image was reoriented to standard orientation (‘RPI’), bias field corrected, and the brain was extracted by the brain extraction tool of FSL. For each subject, the 8 masks were manually checked and edited in order to assure the inclusion of the lower medulla and to discard of noisy non-brain voxel. The corrected mask was then applied to phase images after they were reoriented to standard orientation and converted to angular frequency (radians per second). Then, the angular frequency image was smoothed with a 3D Gaussian kernel with a sigma of 4 mm. The magnitude image was warped to match the angular frequency image (FSL fugue command) and used to linearly register the angular frequency image to the fMRI (FSL flirt command). The angular frequency image registered to the fMRI was used to compute the warp for fMRI distortion correction using the FSL program fugue.

### 7 Tesla fMRI Preprocessing

RETROICOR [Glover et al., 2000] was applied to each run in order to correct for noise related to the phase of physiological cycles, using a custom-built Matlab function able to adapt the RETROICOR algorithm to our slice acquisition order. Functional images were then interpolated in time to align the intra-volume slice timing, and reoriented to standard orientation using FSL routines. Motion correction was applied (FreeSurfer) using the first volume of the first run as reference volume for all the three runs. Six time-series describing rigid motion (three translations and three rotations) of the head were saved for later usage as nuisance regressors. To obtain a reference fMRI volume for the coregistration to the MPRAGE image with enhanced spatial SNR, the first run of the motion-corrected fMRI data was time-averaged (FSL), bias-field corrected (SPM8), and distortion corrected by applying the warp computed (FSL) from the fieldmap. For each subject, the coregistration of the fMRI to the MEMPRAGE was implemented in AFNI (AFNI, Bethesda, MD, USA [Cox, 1996]) using a two-step procedure made of an affine coregistration (AFNI program 3dAllineate) and a nonlinear one. The nonlinear coregistration, implemented in AFNI with the program 3dQwarp, was based on an iterative approach with progressively finer refinement levels and edges enhancement. For 17 subjects out of 20, three refinement levels were used, while for the remaining three subjects results were not satisfactory, thus, an additional level was added. In order to reduce the number of spatial interpolations and the subsequent loss of spatial definition, the affine and nonlinear coregistrations to the MPRAGE were applied to the fMRIs in a single step together with motion correction and distortion correction using AFNI program 3dNwarpApply. After coregistration, a set of nuisance regressors were computed and regressed out in Matlab. Among these nuisance regressors we included the six rigid-body motion time-series, a regressor describing respiratory volume per unit time convolved with a respiration response function [Birn et al., 2008], a regressor describing heart rate convolved with a cardiac response function [Chang et al., 2009] and five regressors modelling the signal in cerebrospinal fluid (CSF).

These latter five CSF regressors were extracted from a principal-components analysis (PCA) performed on a mask of the CSF neighboring the brainstem, defined by thresholding the EPIs standard deviation over time within a 5.3 × 6.5 × 7.5 cm cuboid containing the brainstem. This masks mainly marked voxels in the CSF surrounding the brainstem including the lower portion of the third ventricle, the cerebral aqueduct and fourth ventricle. Previous research proved that the removal of the CSF signal, particularly in the cerebral aqueduct and fourth ventricle, is crucial in the functional connectivity analysis of adjacent nuclei [Satpute et al., 2013; Bianciardi et al., 2016]. Cleaned data were then divided by the temporal signal mean and multiplied by 100 in order to have a timecourse mean of 100, and be allowed to interpret the BOLD signal fluctuations as percentage signal changes with respect to the mean. Bandpass filtering was applied between 0.01 Hz and 0.1 Hz using AFNI program 3dBandpass. Finally, any residual temporal mean was removed and the runs were concatenated.

### Definition of seed and target regions

For the subsequent analyses, a probabilistic atlas of brainstem nuclei in MNI space developed by our team and FreeSurfer’s cortical and subcortical parcellation [Destrieux et al., 2010] were used to define seed and target regions. We used as *seed regions* the probabilistic atlas labels (binarized by setting a threshold at 35%) of the following 15 brainstem nuclei (12 bilateral and 3 midline nuclei, for a total of 27 nuclei) involved in autonomic, limbic, pain and sensory processing: raphe magnus (RMg), lateral parabrachial nucleus (LPB, left and right), medial parabrachial nucleus (MPB, left and right), ventral tegmental area with parabrachial pigmented nucleus of the ventral tegmental area (VTA-PBP, left and right), raphe pallidus (RPa), raphe obscurus (ROb), viscero-sensory-motor complex (VSM, left and right), superior medullary reticular formation (sMRt, left and right), inferior medullary reticular formation (iMRt, left and right), parvicellular reticular nucleus alpha part (PCRtA, left and right), vestibular nuclei complex (Ve, left and right), superior colliculus (SC, left and right), inferior colliculus (IC, left and right) and superior olivary complex (SOC, left and right) and microcellular tegmental nucleus with parabigeminal nucleus (MiTg-PBG, left and right).

We defined as target regions the 27 seed regions, the 164 elements of the FreeSurfer cortical and subcortical parcellation, and 18 (31 counting bilateral nuclei) probabilistic atlas labels (binarized by setting a threshold at 35%) involved in wakeful arousal and motor function (the functional connectome of the latter was computed in [Bianciardi et al., 2016; Singh et al., 2021a]). Masks of brainstem seed and target regions are defined in MNI space. Thus, the average timecourses within brainstem seed and target regions was extracted from the functional images after applying the MEMPRAGE to MNI coregistration transformation described above. FreeSurfer parcellations are defined in single subjects MEMPRAGE space. The average timecourses for FreeSurfer-based target masks were thus extracted from the fMRI data coregistered to MEMPRAGE space. Timecourse extraction from seeds and targets was performed on unsmoothed data.

### Region-based analysis

The Pearson’s correlation coefficient between each seed and target region was computed at the subject level in Matlab using the corrcoeff.m function to yield a correlation matrix. We then applied the Fisher transform (Matlab) to the correlation coefficients to transform the data distribution to be approximately Gaussian (*with null mean and unitary standard deviation*). On the Fisher Z-values arranged in a ‘connectivity’ matrix we could thus implement a two-tailed one-sample t-test to obtain a group statistic. Significance at group level was assessed with a threshold of p<0.0005 Bonferroni corrected for multiple comparisons. Significant connections were displayed for each seed on a circular plot (Matlab code, similar to [Irimia et al., 2012]).

### Voxel-based analysis

To perform voxel-based analysis, fMRI data in MNI space were smoothed with a Gaussian kernel with 1.25mm standard deviation (FWHM=3 mm). For the volumetric analysis, AFNI programs were employed: the timecourse previously extracted from each seed (on *unsmoothed* data) was used as regressor in a general linear model using AFNI program 3dDeconvolve; then, group-level estimates were computed on single-level beta parameters with a two-tailed one sample t-test using AFNI program 3dttest++. False-discovery rate was controlled imposing a lower threshold for activation-cluster size computed with AFNI program 3dClustSim from the estimated autocorrelation function of the residuals. For display purposes of the voxel-based activation maps, we employed a per-voxel threshold of p<0.00001 and a cluster-wise threshold of p<0.01 for nuclei IC, MPB, Ve, iMRt and VSM; for less densely connected nuclei (SC, VTA-PBP, MiTg-PBG, LPB, SOC, sMRt, RMg, RPa, Rob and PCRtA) the less-stringent per-voxel p<0.0001 and cluster-wise p<0.05 thresholds were used for display purposes. We used these thresholds to display activations at the subcortical level on volumetric data, as well as cortical activations, the latter projected on the pial surface using FreeSurfer program mri_vol2surf.

### Diagram generation

We generated a schematic diagram of the autonomic/limbic/nociceptive functional circuit, of the vestibular sensory circuit, and of the interactions between brainstem vestibular nuclei and 11 autonomic/limbic/nociceptive regions. We used as nodes the brainstem nuclei and cortical macroregions involved in each function based on human and animal literature [Balaban, 2004; Indovina et al., 2020; de Lacalle and Saper, 2000; Saper and Stornetta, 2015; Yasui et al., 1989], and as links the connectivity values of our human connectome. Specifically, we averaged out the connectivity strength of subregions belonging to a same node and, for each node, of left and right values, to yield a single connectivity value among nodes. We varied the line thickness of each link based on the connectivity significance, used solid lines for brainstem-to-brain links, and used dashed lines for the remaining (non-brainstem to non-brainstem) links. The same threshold used for the circular plots, i.e. p<0.0005 Bonferroni corrected, was used to assess significance of connections. Note that for the diagram generation, to more precisely define some regions based on their function, we added some newly defined targets in our connectivity matrix, extracted from the FreeSurfer parcellation [Destrieux et al., 2010] and from the Eickhoff-Fan atlas [Eickhoff et al., 2005; Fan et al., 2016]. Specifically, in the autonomic/limbic/pain circuit, we added the: (1) Infralimbic Cortex, defined as the macroregion containing rectus gyrus and subcallosal gyrus from FreeSurfer parcellation; (2) Orbitofrontal Cortex, including the orbital part of the inferior frontal gyrus, orbital gyrus, horizontal ramus and vertical ramus of the anterior segment of the lateral sulcus, lateral orbital sulcus, medial orbital (olfactory) sulcus, orbital h-shaped sulcus and suborbital sulcus from FreeSurfer parcellation; (3) Anterior Cingulate Cortex with anterior and middle-anterior parts of the cingulate gyrus and sulcus form FreeSurfer parcellation; (4) Anterior Insula, including dorsal agranular insula, ventral agranular insula and temporal agranular insular cortex from Eickhoff-Fan parcellation; (5) Fastigii,Nucleus from Eickhoff-Fan parcellation. In the vestibular circuit, we added the: (5) Fastigii,Nucleus from Eickhoff-Fan parcellation; (6) cerebellar Lobule X from Eickhoff-Fan parcellation; (7) Intraparietal Sulcus, including areas hIP1, hIP2 and hIP3 from Eickhoff-Fan parcellation; (8) Temporal Sulcus, with rostroposterior and caudoposterior temporal sulcus from Eickhoff-Fan parcellation; (9) Posterior Cingulate Cortex, including dorsal, caudal and ventral area 23 and caudodorsal area 24 from Eickhoff-Fan parcellation plus marginal branch of the cingulate sulcus from FreeSurfer parcellation; (10) Inferior Frontal Gyrus, including area 44 with its dorsal, opercular and ventral parts, and area 45 with its caudal part from Eichoff-Fan parcellation; (11) Parietal Operculum, with areas OP1, OP2, OP3 and OP4 from Eickhoff-Fan parcellation; (12) Posterior Insula, including area Ig1, Ig2, hypergranular insula, dorsal granular insula and ventral dysgranular and granular insula from Eickhoff-Fan parcellation; (13) Middle Insula, including area Id1 and dorsal dysgranular insula from Eickhoff-Fan parcellation; (14) Pre-Motor Cortex, including caudal dorsolateral and caudal ventrolateral area 6 from Eickhoff-Fan parcellation; (15) Superior Parietal Lobe, with intraparietal and postcentral area 7, area 7A, lateral area 5, as well as area 5Ci from Eickhoff-Fan parcellation; (16) Precuneus, including dorsomedial parieto-occipital sulcus, regions 5M, 7M, medial area 7, medial area 5, and area 31 from Eickhoff-Fan parcellation; (17) Inferior Parietal Cortex, including rostroventral, rostrodorsal and caudal area 39, rostroventral, rostrodorsal and caudal area 40, area Pfm, area PF, area PFcm, area PFt, area PGp from Eickhoff-Fan parcellation; (18) Visual-Motion Cortex, including regions V5/MT+ and hOC5 from Eickhoff-Fan parcellation; (19) Fusiform Cortex including dorsolateral, ventrolateral, medioventral and lateroventral area 37 from Eickhoff-Fan parcellation. For the calculation of the Bonferroni corrected statistical threshold, the degrees of freedom were corrected to include these additional 19 regions.

### 3 Tesla fMRI Preprocessing

fMRI data acquired on the 3 Tesla scanner underwent a preprocessing pipeline as similar as possible to the one used with 7 Tesla functional data. The only relevant difference was in the parameters used for the nonlinear coregistration between fMRI and MEMPRAGE: after visual assessment of coregistration quality for various sets of parameters, for 18 subjects we employed the same set used for the majority of 7 Tesla images, but with no edge enhancement. For the remaining 2 subjects, we opted to keep the only affine step of coregistration, with no additional nonlinear warp.

The replicability of 7 Tesla results on 3 Tesla data was assessed by computing the correlation coefficient between the two region-based correlation matrices (3 Tesla vs 7 Tesla), averaged across subjects. Significance of the correlation was assessed in all cases at p < 0.05.

## Results

In Figure 1A, we show the quality of the coregistration between functional images and the MNI template, on a mid-sagittal slice zoomed on the brainstem as shown in Figure 1B. In addition, in Figure 1C we show the position of brainstem seeds and targets in the MNI space to prove that none of them is located in anterior regions that might be affected by residual distortions or signal dropout. In Figure 1D we display the effect of distortion correction on the functional image of subject 1.

**Figure 1.**
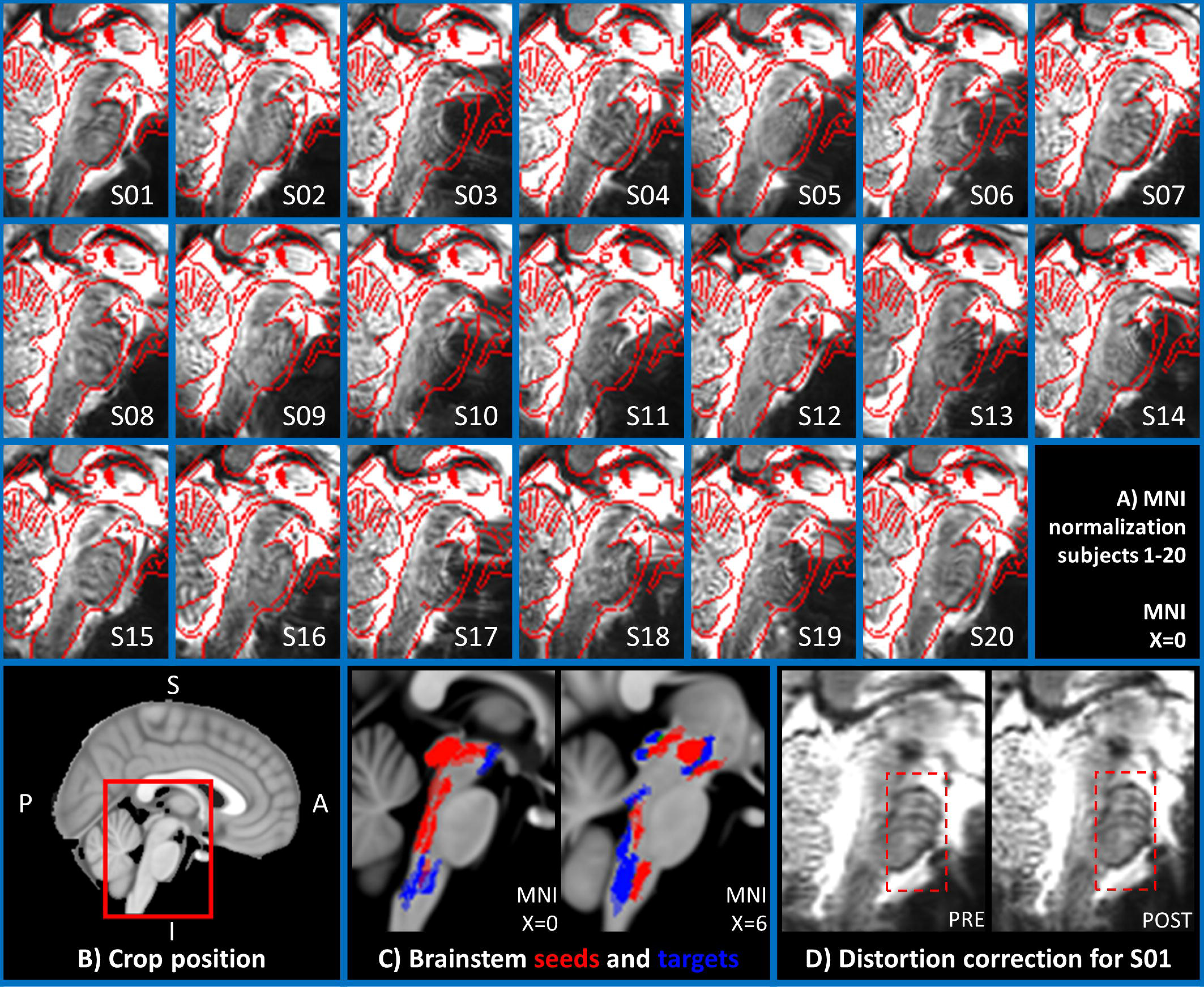
Quality of the fMRI coregistration to MNI space: In the 20 images composing panel A, the quality of the coregistration is evaluated as follows: In the underlay, we display the time averaged motion corrected fMRI of the first run, which was used as reference image to compute the coregistration transformation to the subject’s MEMPRAGE. The edges of the MNI standard template are overlaid in red for quality control. Images were generated for all subjects (S1-20) using FSL slices on the most medial sagittal slice (MNI space, x=0). In B) we clarify on a midsagittal view of the MNI template the cropping and orientation used for the images in panel A. In C) we show on two sagittal slices of the MNI template the location of all seeds (in red) and all brainstem targets (in blue). In D) we show for subject 01 the time averaged motion corrected fMRI of the first run, before and after applying distortion correction. The pons, which is more prone to distortions, is highlighted with a red box.

In Figure 2 we report the seeds-to-brain (27-by-227) matrix of p-values for group-level correlation values, after taking the opposite of their base-10 logarithm. A Bonferroni-corrected threshold p<0.0005 was used for display purposes. Overall density at this threshold was 68%. As visible from the matrix, seeds-to-others connectivity was denser than the seeds-to-brainstem one (73% vs 56% density). The average correlation coefficient was generally higher (p<0.001, two-samples t-test) for brainstem-to-cortex links (mean±std: 0.22±0.08) with respect to brainstem-to-brainstem ones (mean±std: 0.14±0.07), but brainstem-to-brainstem links (mean±std: 0.071±0.016) displayed lower (p<0.001, two-sample t-test) standard deviations from the average than brainstem-to-cortex links (mean±std: 0.094±0.023), see also Supplementary Figures S1, displaying the circular plot of all the seeds-to-brain significant functional connections (p<0.0005, Bonferroni corrected) and S2, S3 and S4 which report in matrix form the average connectivity values, their standard deviation, and the opposite of the base-10 logarithm of p values without thresholds. In Figures S2, S3 and S4 we also include cortex-to-cortex results.

**Figure 2.**
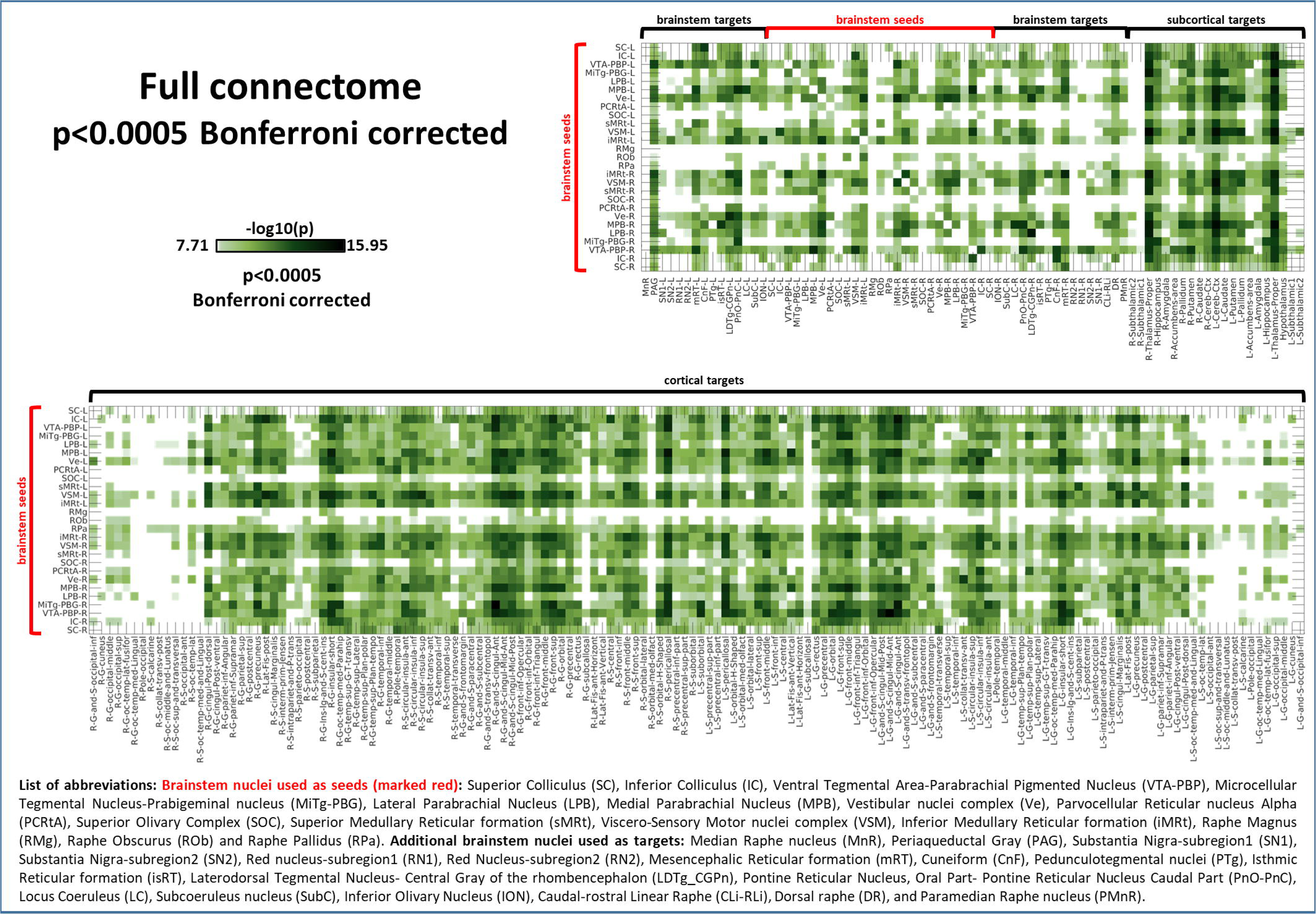
Seeds-to-Brain correlation matrix displaying the –log_10_(p-value) extracted from the group-level region-based analysis of seeds-to-brain functional connectivity (p<0.0005, Bonferroni corrected). **List of abbreviations:** Brainstem nuclei used as seeds (marked with red brackets): Superior Colliculus (SC), Inferior Colliculus (IC), Ventral Tegmental Area-Parabrachial Pigmented Nucleus (VTA-PBP), Microcellular Tegmental Nucleus-Prabigeminal nucleus (MiTg-PBG), Lateral Parabrachial Nucleus (LPB), Medial Parabrachial Nucleus (MPB), Vestibular nuclei complex (Ve), Parvicellular Reticular nucleus Alpha (PCRtA), Superior Olivary Complex (SOC), Superior Medullary Reticular formation (sMRt), Viscero-Sensory Motor nuclei complex (VSM), Inferior Medullary Reticular formation (iMRt), Raphe Magnus (RMg), Raphe Obscurus (ROb) and Raphe Pallidus (RPa). Additional brainstem nuclei used as targets: Median Raphe nucleus (MnR), Periaqueductal Gray (PAG), Substantia Nigra-subregion1 (SN1), Substantia Nigra-subregion2 (SN2), Red nucleus-subregion1 (RN1), Red Nucleus-subregion2 (RN2), Mesencephalic Reticular formation (mRt), Cuneiform (CnF), Pedunculotegmental nuclei (PTg), Isthmic Reticular formation (isRt), Laterodorsal Tegmental Nucleus-Central Gray of the rhombencephalon (LDTg_CGPn), Pontine Reticular Nucleus, Oral Part-Pontine Reticular Nucleus Caudal Part (PnO-PnC), Locus Coeruleus (LC), Subcoeruleus nucleus (SubC), Inferior Olivary Nucleus (ION), Caudal-rostral Linear Raphe (CLi-RLi), Dorsal raphe (DR), and Paramedian Raphe nucleus (PMnR).

We computed the laterality index for each bilateral seed: values are as follows: SC 15.3%, IC 3.6%, VTA-PBP 3.7%, MiTg-PBG 10.3%, LPB 6.7%, MPB 6.3%, Ve 1.3%, PCRtA 12.8%, SOC 17.7%, sMRt 5.4%, VSM 2.2%, iMRt 1.3%. No seed exceeded 18% laterality, with only SOC, SC, PCRtA and MiTg-PBG exceeding 10%. The values are reported graphically in Figure S5, panel C in the supplementary materials.

In Figures 3-10, we report the functional connectivity computed between 27 brainstem seeds (which we refer to as “seeds”) and 227 targets (which we call “brain”), 58 located within the brainstem (which we refer to as “brainstem”), and 169 in cortical and subcortical regions (which we call “rest of the brain” or “others”). We refer to the 148 cortical targets as “cortex”.

In Figures 3-10 detailed functional connectivity results are reported separately for each seed (Figure 3: SC (top), IC (bottom); Figure 4: VTA-PBP (top), MiTg-PBG (bottom); Figure 5: LPB (top), MPB (bottom); Figure 6: Ve (top), SOC (bottom); Figure 7: sMRt (top), iMRt (bottom); Figure 8: VSM (top), RMg (bottom); Figure 9: RPa (top), ROb (bottom); Figure 10: PCRtA) in the form of a region-based circular connectome (panels A, as in Figure 2 and at the same significance threshold p<0.0005, Bonferroni-corrected). In addition, for each seed we show the voxel-based functional connectivity maps displayed with FreeSurfer on the medial and lateral cortex of both hemispheres (panels B), and subcortically with FSLview for three orthogonal views (panels C).

**Figure 3.**
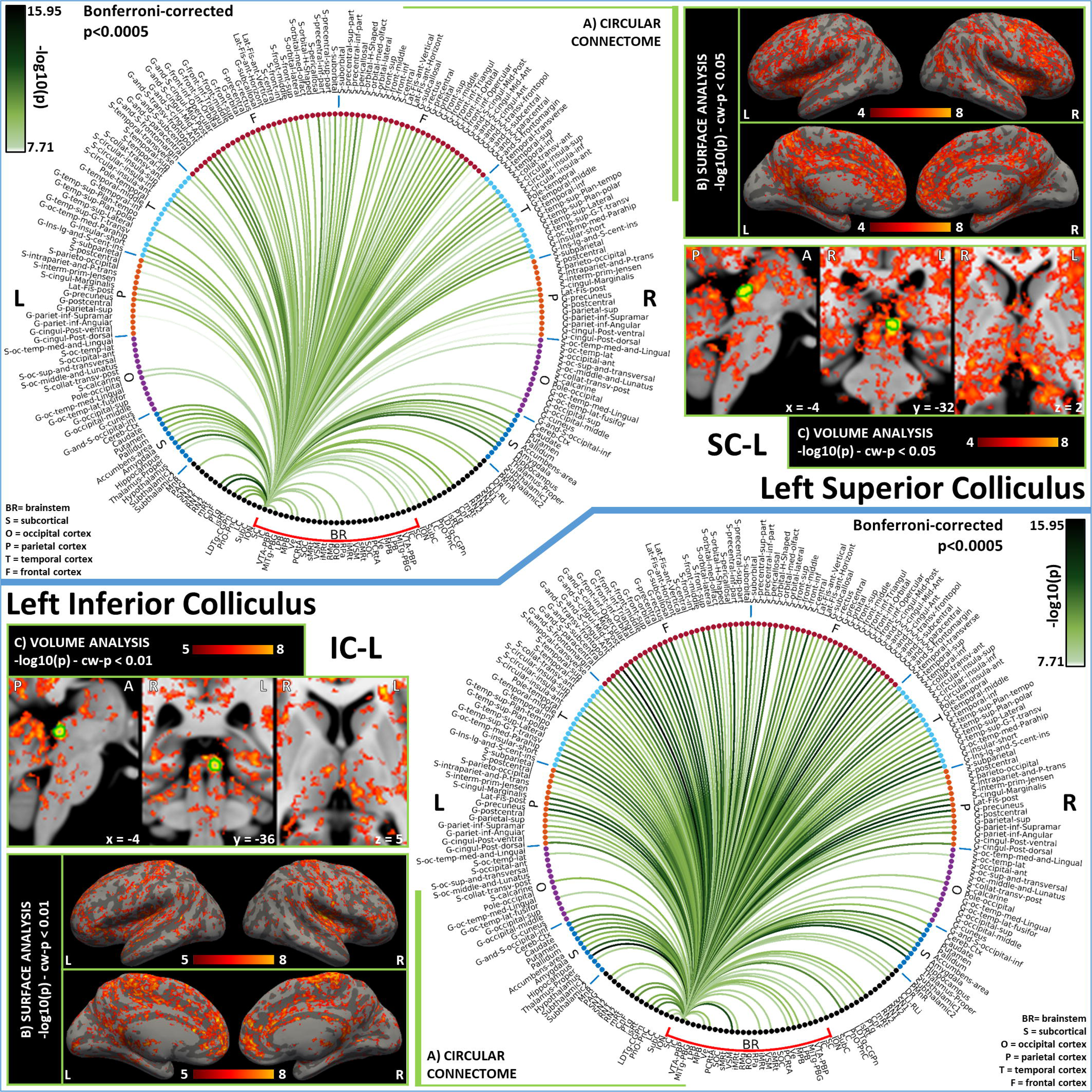
Functional connectivity results of Superior Colliculus – SC (top) and Inferior Colliculus - IC (bottom). **A)** Region-based functional connectome; **B)** voxel-based cortical connectivity maps displayed on two lateral and two medial surface views; **C)** voxel-based subcortical connectivity maps, three views, MNI coordinates. The seed is highlighted with green contours. **List of abbreviations:** Brainstem nuclei used as seeds (marked with a red bracket): Superior Colliculus (SC), Inferior Colliculus (IC), Ventral Tegmental Area-Parabrachial Pigmented Nucleus (VTA-PBP), Microcellular Tegmental Nucleus-Prabigeminal nucleus (MiTg-PBG), Lateral Parabrachial Nucleus (LPB), Medial Parabrachial Nucleus (MPB), Vestibular nuclei complex (Ve), Parvicellular Reticular nucleus Alpha (PCRtA), Superior Olivary Complex (SOC), Superior Medullary Reticular formation (sMRt), Viscero-Sensory Motor nuclei complex (VSM), Inferior Medullary Reticular formation (iMRt), Raphe Magnus (RMg), Raphe Obscurus (ROb) and Raphe Pallidus (RPa). Additional brainstem nuclei used as targets: Median Raphe nucleus (MnR), Periaqueductal Gray (PAG), Substantia Nigra-subregion1 (SN1), Substantia Nigra-subregion2 (SN2), Red nucleus-subregion1 (RN1), Red Nucleus-subregion2 (RN2), Mesencephalic Reticular formation (mRT), Cuneiform (CnF), Pedunculotegmental nuclei (PTg), Isthmic Reticular formation (isRt), Laterodorsal Tegmental Nucleus-Central Gray of the rhombencephalon (LDTg_CGPn), Pontine Reticular Nucleus, Oral Part-Pontine Reticular Nucleus Caudal Part (PnO-PnC), Locus Coeruleus (LC), Subcoeruleus nucleus (SubC), Inferior Olivary Nucleus (ION), Caudal-rostral Linear Raphe (CLi-RLi), Dorsal raphe (DR), and Paramedian Raphe nucleus (PMnR).

**Figure 4.**
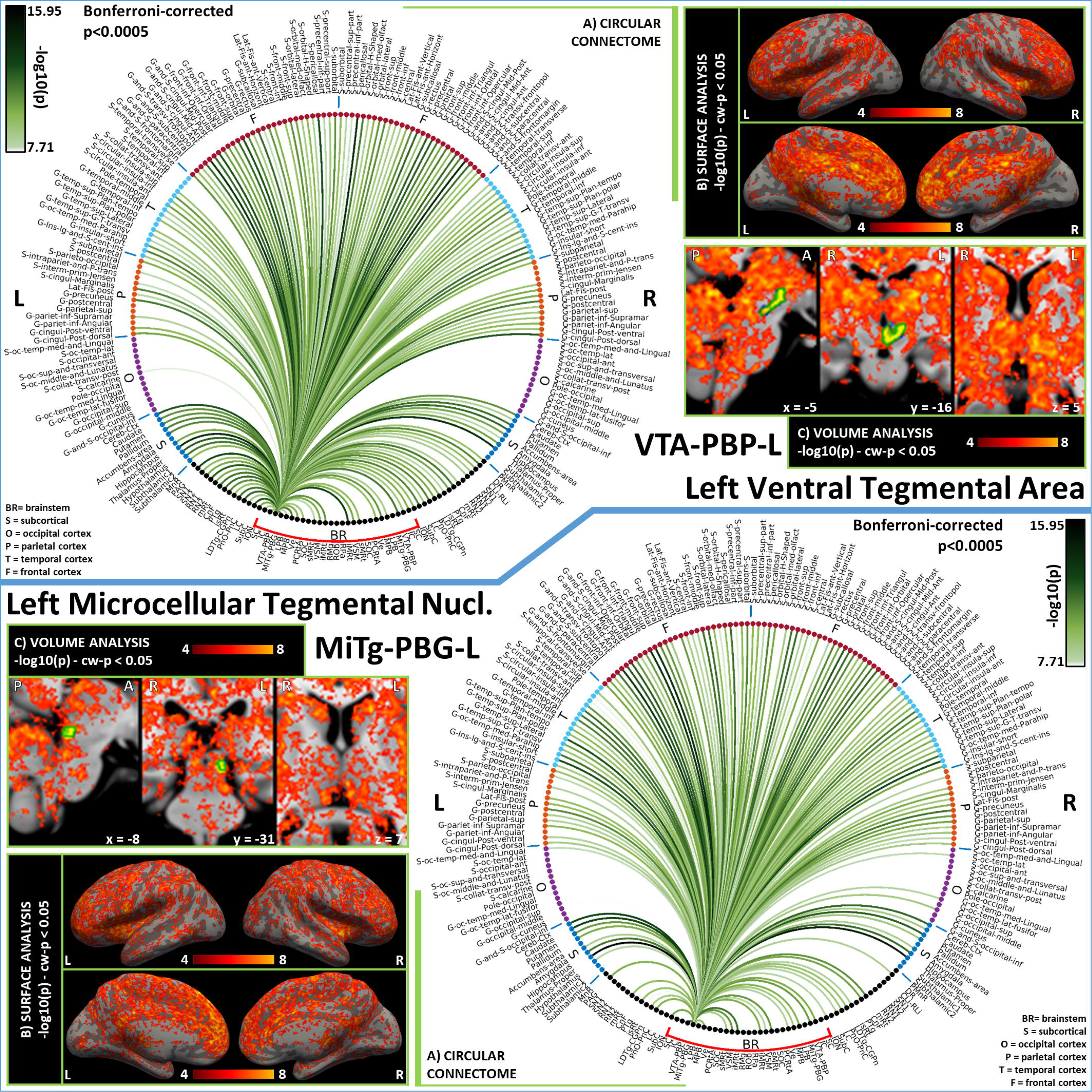
Functional connectivity results of Left Ventral Tegmental Area – VTA-PBP (top) and Left Microcellular Tegmental Nucelus – MiTg-PBG (bottom). **A)** Region-based functional connectome; **B)** voxel-based cortical connectivity maps displayed on two lateral and two medial surface views; **C)** voxel-based subcortical connectivity maps, three views, MNI coordinates. The seed is highlighted with green contours. **List of abbreviations:** Brainstem nuclei used as seeds (marked with a red bracket): Superior Colliculus (SC), Inferior Colliculus (IC), Ventral Tegmental Area-Parabrachial Pigmented Nucleus (VTA-PBP), Microcellular Tegmental Nucleus-Prabigeminal nucleus (MiTg-PBG), Lateral Parabrachial Nucleus (LPB), Medial Parabrachial Nucleus (MPB), Vestibular nuclei complex (Ve), Parvicellular Reticular nucleus Alpha (PCRtA), Superior Olivary Complex (SOC), Superior Medullary Reticular formation (sMRt), Viscero-Sensory Motor nuclei complex (VSM), Inferior Medullary Reticular formation (iMRt), Raphe Magnus (RMg), Raphe Obscurus (ROb) and Raphe Pallidus (RPa). Additional brainstem nuclei used as targets: Median Raphe nucleus (MnR), Periaqueductal Gray (PAG), Substantia Nigra-subregion1 (SN1), Substantia Nigra-subregion2 (SN2), Red nucleus-subregion1 (RN1), Red Nucleus-subregion2 (RN2), Mesencephalic Reticular formation (mRt), Cuneiform (CnF), Pedunculotegmental nuclei (PTg), Isthmic Reticular formation (isRt), Laterodorsal Tegmental Nucleus-Central Gray of the rhombencephalon (LDTg_CGPn), Pontine Reticular Nucleus, Oral Part-Pontine Reticular Nucleus Caudal Part (PnO-PnC), Locus Coeruleus (LC), Subcoeruleus nucleus (SubC), Inferior Olivary Nucleus (ION), Caudal-rostral Linear Raphe (CLi-RLi), Dorsal raphe (DR), and Paramedian Raphe nucleus (PMnR).

**Figure 5.**
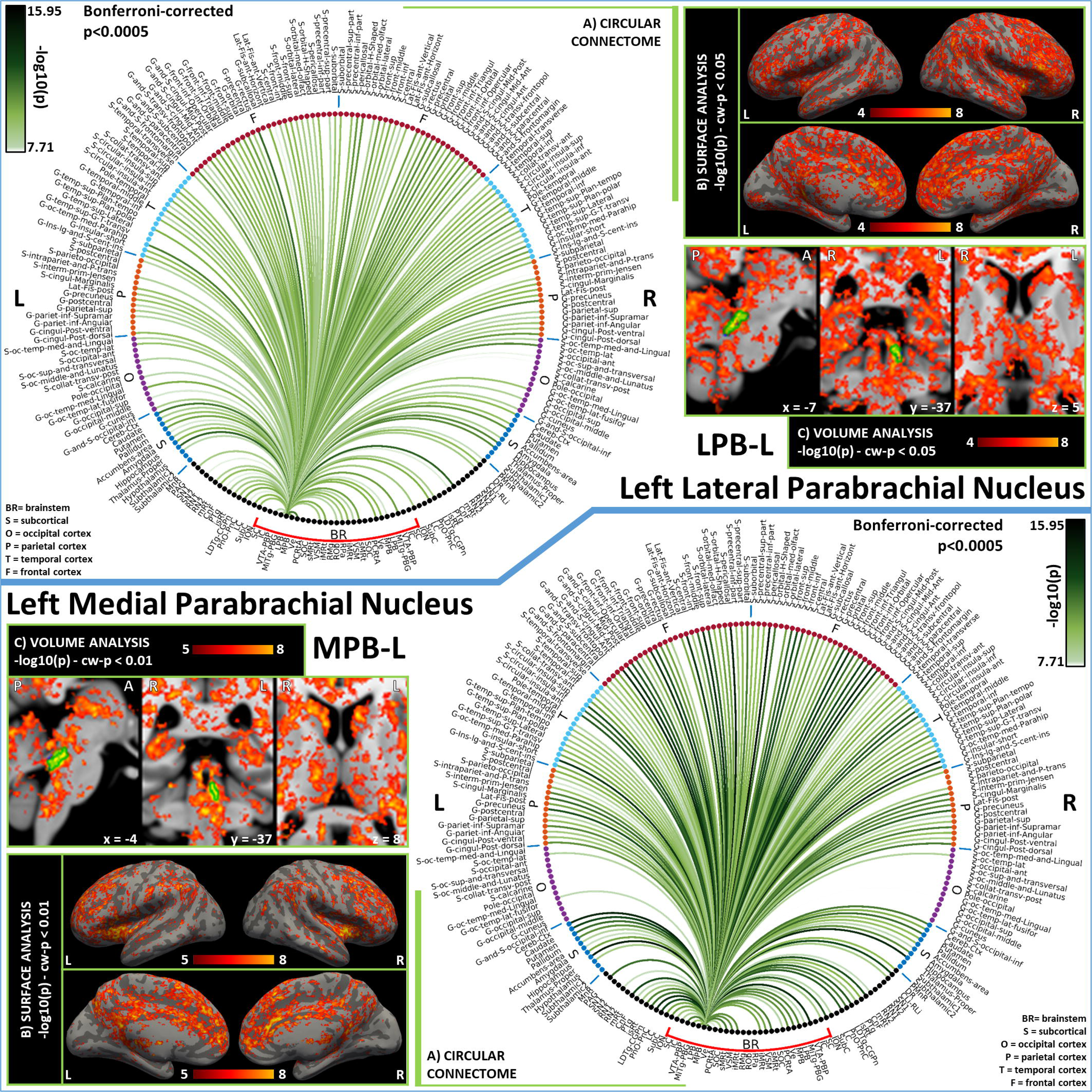
Functional connectivity results of Left Lateral Parabrachial Nucleus – LPB (top) and Left Medial Parabrachial Nucleus – MPB (bottom). **A)** Region-based functional connectome; **B)** voxel-based cortical connectivity maps displayed on two lateral and two medial surface views; **C)** voxel-based subcortical connectivity maps, three views, MNI coordinates. The seed is highlighted with green contours. **List of abbreviations:** Brainstem nuclei used as seeds (marked with a red bracket): Superior Colliculus (SC), Inferior Colliculus (IC), Ventral Tegmental Area-Parabrachial Pigmented Nucleus (VTA-PBP), Microcellular Tegmental Nucleus-Prabigeminal nucleus (MiTg-PBG), Lateral Parabrachial Nucleus (LPB), Medial Parabrachial Nucleus (MPB), Vestibular nuclei complex (Ve), Parvicellular Reticular nucleus Alpha (PCRtA), Superior Olivary Complex (SOC), Superior Medullary Reticular formation (sMRt), Viscero-Sensory Motor nuclei complex (VSM), Inferior Medullary Reticular formation (iMRt), Raphe Magnus (RMg), Raphe Obscurus (ROb) and Raphe Pallidus (RPa). Additional brainstem nuclei used as targets: Median Raphe nucleus (MnR), Periaqueductal Gray (PAG), Substantia Nigra-subregion1 (SN1), Substantia Nigra-subregion2 (SN2), Red nucleus-subregion1 (RN1), Red Nucleus-subregion2 (RN2), Mesencephalic Reticular formation (mRt), Cuneiform (CnF), Pedunculotegmental nuclei (PTg), Isthmic Reticular formation (isRt), Laterodorsal Tegmental Nucleus-Central Gray of the rhombencephalon (LDTg_CGPn), Pontine Reticular Nucleus, Oral Part-Pontine Reticular Nucleus Caudal Part (PnO-PnC), Locus Coeruleus (LC), Subcoeruleus nucleus (SubC), Inferior Olivary Nucleus (ION), Caudal-rostral Linear Raphe (CLi-RLi), Dorsal raphe (DR), and Paramedian Raphe nucleus (PMnR).

**Figure 6.**
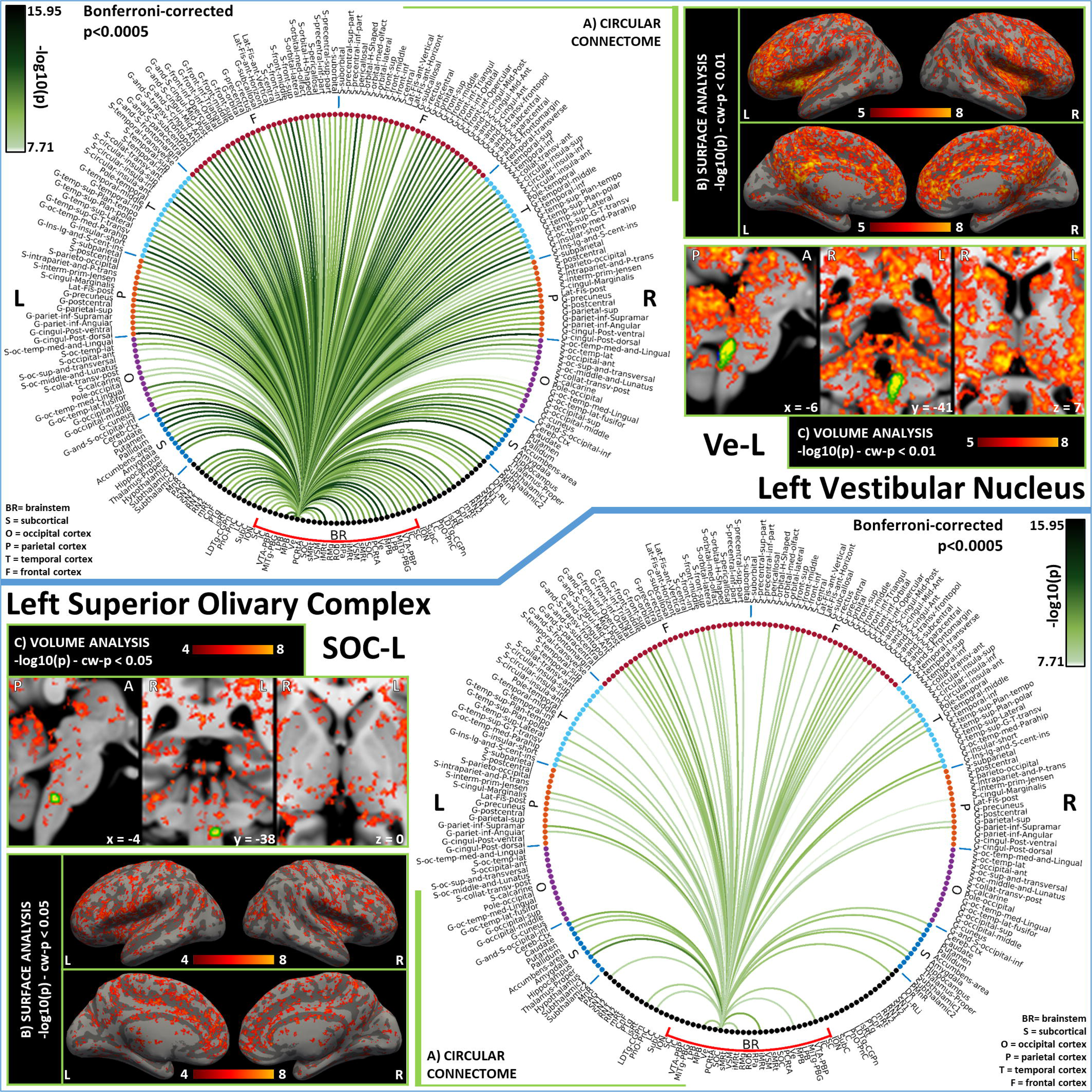
Functional connectivity results of Left Vestibular Nucleus – Ve (top) and Left Superior Olivary Complex – SOC (bottom). **A)** Region-based functional connectome; **B)** voxel-based cortical connectivity maps displayed on two lateral and two medial surface views; **C)** voxel-based subcortical connectivity maps, three views, MNI coordinates. The seed is highlighted with green contours. **List of abbreviations:** Brainstem nuclei used as seeds (marked with a red bracket): Superior Colliculus (SC), Inferior Colliculus (IC), Ventral Tegmental Area-Parabrachial Pigmented Nucleus (VTA-PBP), Microcellular Tegmental Nucleus-Prabigeminal nucleus (MiTg-PBG), Lateral Parabrachial Nucleus (LPB), Medial Parabrachial Nucleus (MPB), Vestibular nuclei complex (Ve), Parvicellular Reticular nucleus Alpha (PCRtA), Superior Olivary Complex (SOC), Superior Medullary Reticular formation (sMRt), Viscero-Sensory Motor nuclei complex (VSM), Inferior Medullary Reticular formation (iMRt), Raphe Magnus (RMg), Raphe Obscurus (ROb) and Raphe Pallidus (RPa). Additional brainstem nuclei used as targets: Median Raphe nucleus (MnR), Periaqueductal Gray (PAG), Substantia Nigra-subregion1 (SN1), Substantia Nigra-subregion2 (SN2), Red nucleus-subregion1 (RN1), Red Nucleus-subregion2 (RN2), Mesencephalic Reticular formation (mRt), Cuneiform (CnF), Pedunculotegmental nuclei (PTg), Isthmic Reticular formation (isRt), Laterodorsal Tegmental Nucleus-Central Gray of the rhombencephalon (LDTg_CGPn), Pontine Reticular Nucleus, Oral Part-Pontine Reticular Nucleus Caudal Part (PnO-PnC), Locus Coeruleus (LC), Subcoeruleus nucleus (SubC), Inferior Olivary Nucleus (ION), Caudal-rostral Linear Raphe (CLi-RLi), Dorsal raphe (DR), and Paramedian Raphe nucleus (PMnR).

**Figure 7.**
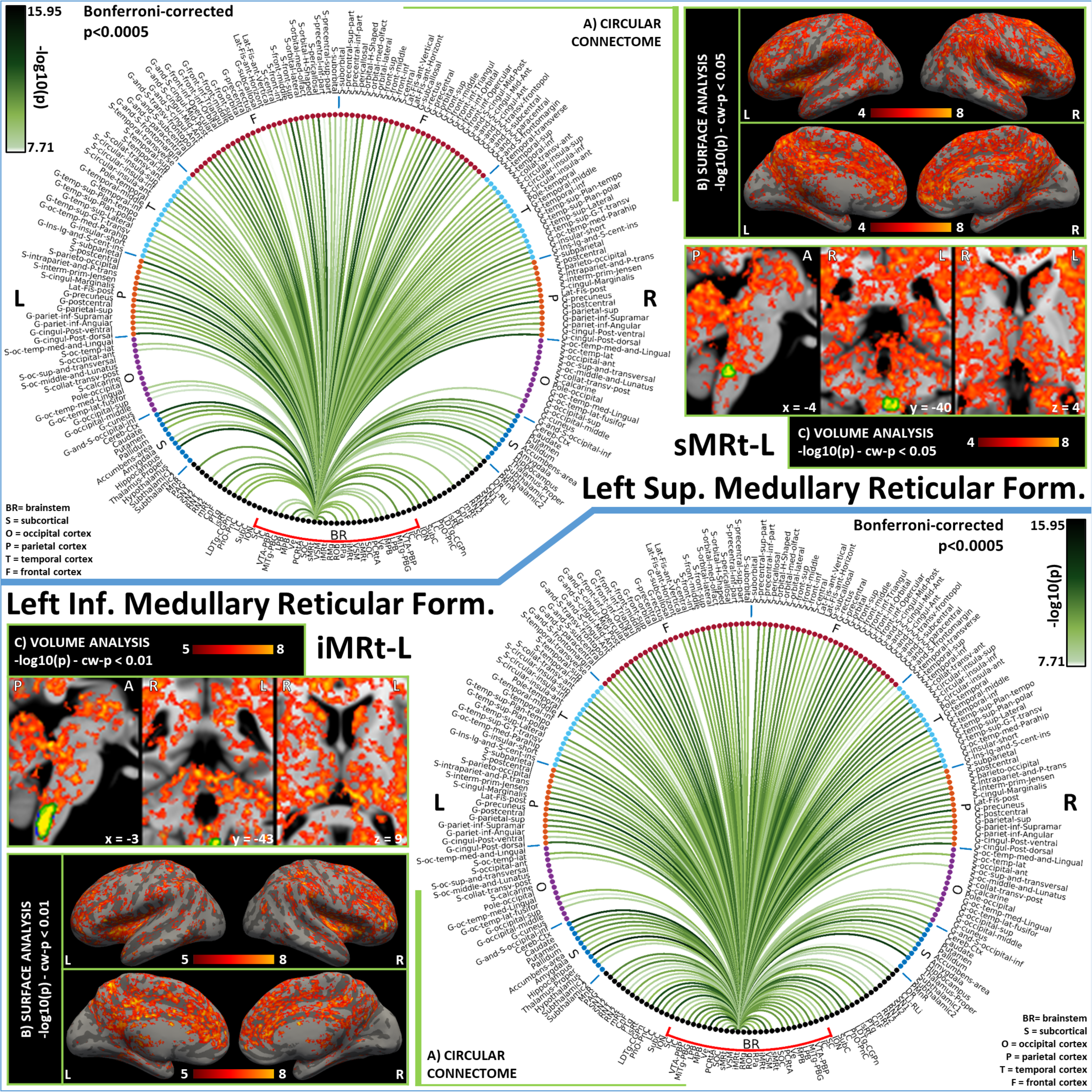
Functional connectivity results of Left Superior Medullary Reticular Formation – sMRt (top) and Left Inferior Medullary Reticular Formation – iMRt (bottom). **A)** Region-based functional connectome; **B)** voxel-based cortical connectivity maps displayed on two lateral and two medial surface views; **C)** voxel-based subcortical connectivity maps, three views, MNI coordinates. The seed is highlighted with green contours. **List of abbreviations:** Brainstem nuclei used as seeds (marked with a red bracket): Superior Colliculus (SC), Inferior Colliculus (IC), Ventral Tegmental Area-Parabrachial Pigmented Nucleus (VTA-PBP), Microcellular Tegmental Nucleus-Prabigeminal nucleus (MiTg-PBG), Lateral Parabrachial Nucleus (LPB), Medial Parabrachial Nucleus (MPB), Vestibular nuclei complex (Ve), Parvicellular Reticular nucleus Alpha (PCRtA), Superior Olivary Complex (SOC), Superior Medullary Reticular formation (sMRt), Viscero-Sensory Motor nuclei complex (VSM), Inferior Medullary Reticular formation (iMRt), Raphe Magnus (RMg), Raphe Obscurus (ROb) and Raphe Pallidus (RPa). Additional brainstem nuclei used as targets: Median Raphe nucleus (MnR), Periaqueductal Gray (PAG), Substantia Nigra-subregion1 (SN1), Substantia Nigra-subregion2 (SN2), Red nucleus-subregion1 (RN1), Red Nucleus-subregion2 (RN2), Mesencephalic Reticular formation (mRt), Cuneiform (CnF), Pedunculotegmental nuclei (PTg), Isthmic Reticular formation (isRt), Laterodorsal Tegmental Nucleus-Central Gray of the rhombencephalon (LDTg_CGPn), Pontine Reticular Nucleus, Oral Part-Pontine Reticular Nucleus Caudal Part (PnO-PnC), Locus Coeruleus (LC), Subcoeruleus nucleus (SubC), Inferior Olivary Nucleus (ION), Caudal-rostral Linear Raphe (CLi-RLi), Dorsal raphe (DR), and Paramedian Raphe nucleus (PMnR).

**Figure 8.**
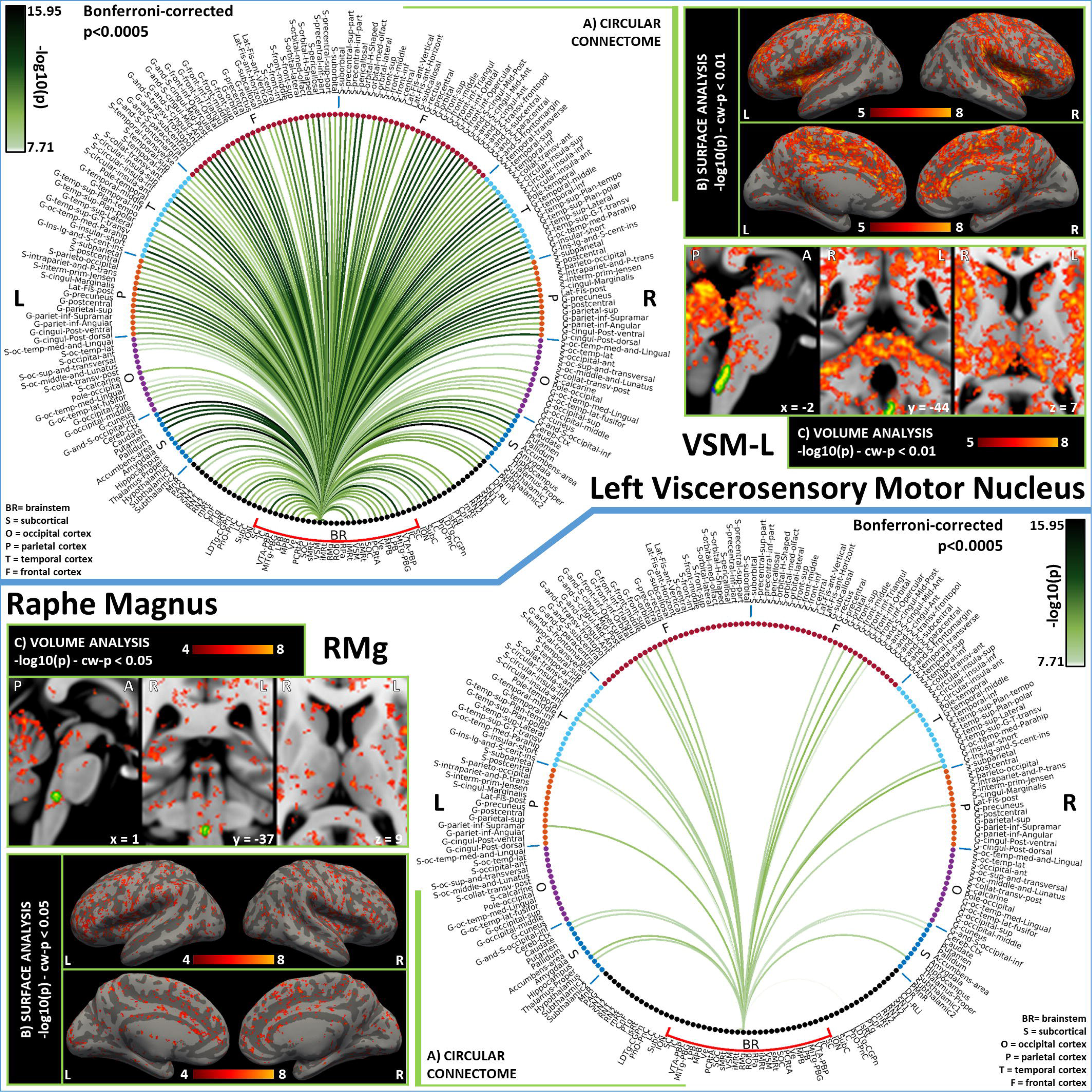
Functional connectivity results of Left Viscerosensory Motor Nucleus – VSM (top) and Raphe Magnus – RMg (bottom). **A)** Region-based functional connectome; **B)** voxel-based cortical connectivity maps displayed on two lateral and two medial surface views; **C)** voxel-based subcortical connectivity maps, three views, MNI coordinates. The seed is highlighted with green contours. **List of abbreviations:** Brainstem nuclei used as seeds (marked with a red bracket): Superior Colliculus (SC), Inferior Colliculus (IC), Ventral Tegmental Area-Parabrachial Pigmented Nucleus (VTA-PBP), Microcellular Tegmental Nucleus-Prabigeminal nucleus (MiTg-PBG), Lateral Parabrachial Nucleus (LPB), Medial Parabrachial Nucleus (MPB), Vestibular nuclei complex (Ve), Parvicellular Reticular nucleus Alpha (PCRtA), Superior Olivary Complex (SOC), Superior Medullary Reticular formation (sMRt), Viscero-Sensory Motor nuclei complex (VSM), Inferior Medullary Reticular formation (iMRt), Raphe Magnus (RMg), Raphe Obscurus (ROb) and Raphe Pallidus (RPa). Additional brainstem nuclei used as targets: Median Raphe nucleus (MnR), Periaqueductal Gray (PAG), Substantia Nigra-subregion1 (SN1), Substantia Nigra-subregion2 (SN2), Red nucleus-subregion1 (RN1), Red Nucleus-subregion2 (RN2), Mesencephalic Reticular formation (mRt), Cuneiform (CnF), Pedunculotegmental nuclei (PTg), Isthmic Reticular formation (isRt), Laterodorsal Tegmental Nucleus-Central Gray of the rhombencephalon (LDTg_CGPn), Pontine Reticular Nucleus, Oral Part-Pontine Reticular Nucleus Caudal Part (PnO-PnC), Locus Coeruleus (LC), Subcoeruleus nucleus (SubC), Inferior Olivary Nucleus (ION), Caudal-rostral Linear Raphe (CLi-RLi), Dorsal raphe (DR), and Paramedian Raphe nucleus (PMnR).

**Figure 9.**
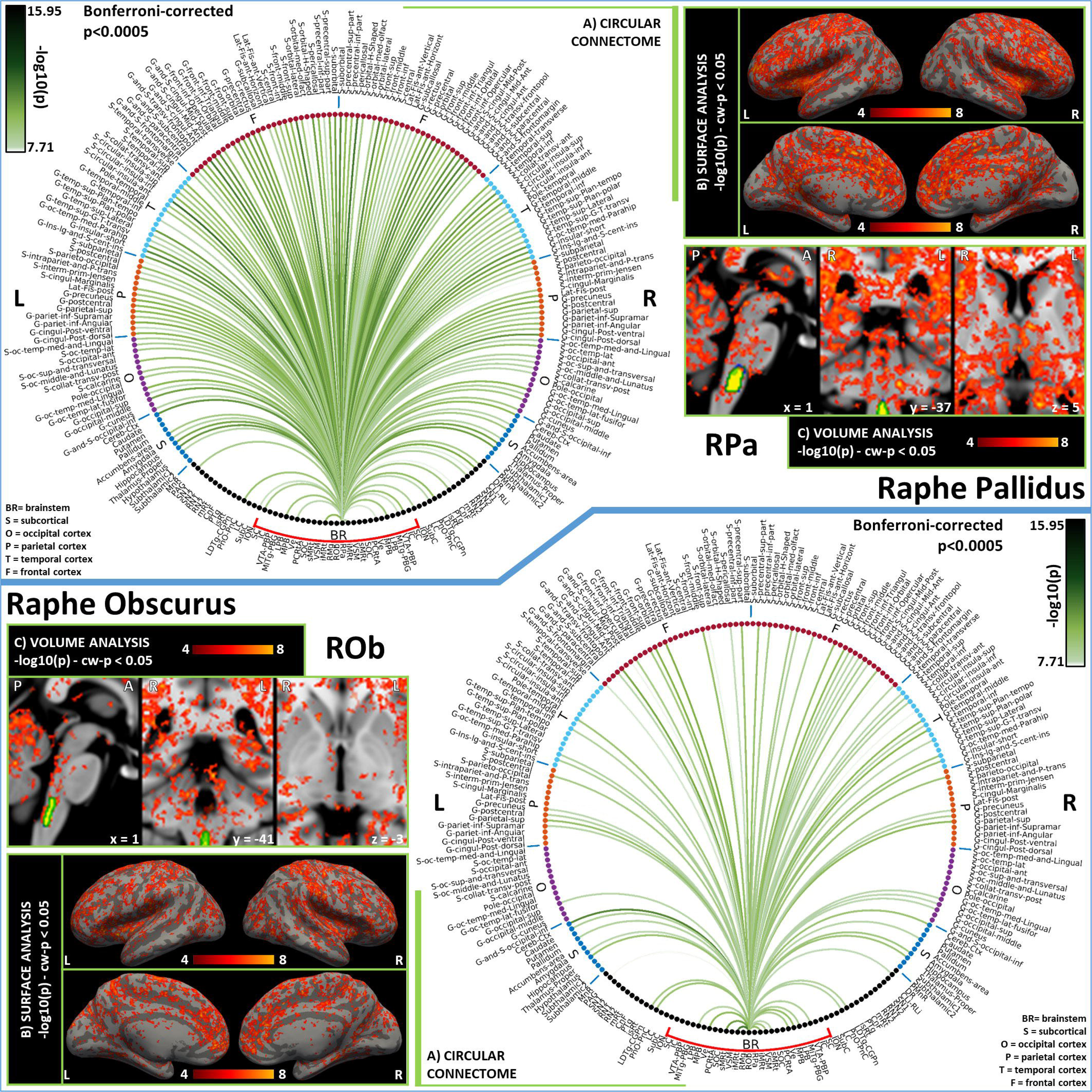
Functional connectivity results of Raphe Pallidus – RPa (top) and Raphe Obscurus – ROb (bottom). **A)** Region-based functional connectome; **B)** voxel-based cortical connectivity maps displayed on two lateral and two medial surface views; **C)** voxel-based subcortical connectivity maps, three views, MNI coordinates. The seed is highlighted with green contours. **List of abbreviations:** Brainstem nuclei used as seeds (marked with a red bracket): Superior Colliculus (SC), Inferior Colliculus (IC), Ventral Tegmental Area-Parabrachial Pigmented Nucleus (VTA-PBP), Microcellular Tegmental Nucleus-Prabigeminal nucleus (MiTg-PBG), Lateral Parabrachial Nucleus (LPB), Medial Parabrachial Nucleus (MPB), Vestibular nuclei complex (Ve), Parvicellular Reticular nucleus Alpha (PCRtA), Superior Olivary Complex (SOC), Superior Medullary Reticular formation (sMRt), Viscero-Sensory Motor nuclei complex (VSM), Inferior Medullary Reticular formation (iMRt), Raphe Magnus (RMg), Raphe Obscurus (ROb) and Raphe Pallidus (RPa). Additional brainstem nuclei used as targets: Median Raphe nucleus (MnR), Periaqueductal Gray (PAG), Substantia Nigra-subregion1 (SN1), Substantia Nigra-subregion2 (SN2), Red nucleus-subregion1 (RN1), Red Nucleus-subregion2 (RN2), Mesencephalic Reticular formation (mRt), Cuneiform (CnF), Pedunculotegmental nuclei (PTg), Isthmic Reticular formation (isRt), Laterodorsal Tegmental Nucleus-Central Gray of the rhombencephalon (LDTg_CGPn), Pontine Reticular Nucleus, Oral Part-Pontine Reticular Nucleus Caudal Part (PnO-PnC), Locus Coeruleus (LC), Subcoeruleus nucleus (SubC), Inferior Olivary Nucleus (ION), Caudal-rostral Linear Raphe (CLi-RLi), Dorsal raphe (DR), and Paramedian Raphe nucleus (PMnR).

**Figure 10.**
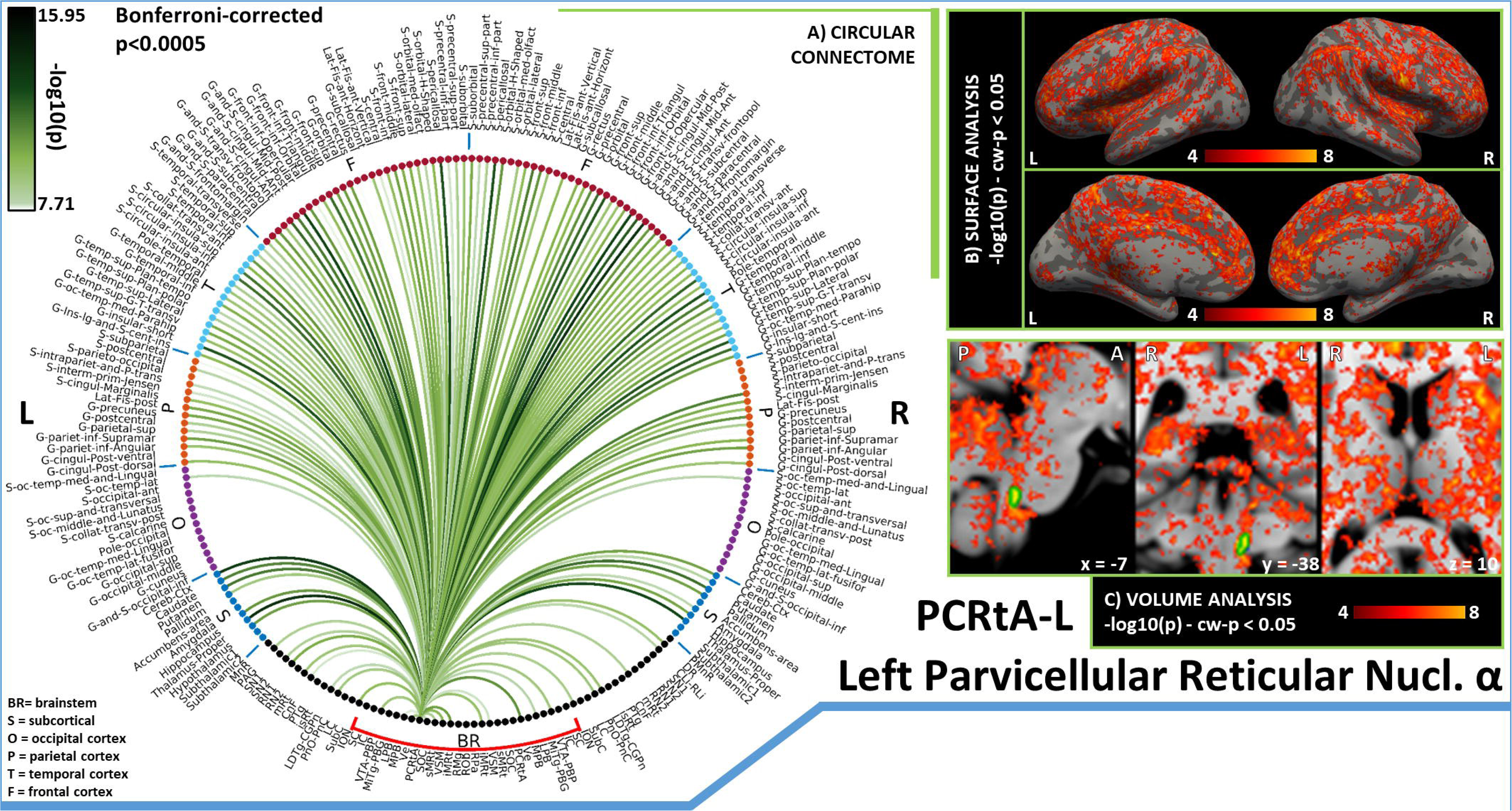
Functional connectivity results of Left Alpha Part of the Parvicellular Reticular Nucleus – PCRtA. **A)** Region-based functional connectome; **B)** voxel-based cortical connectivity maps displayed on two lateral and two medial surface views; **C)** voxel-based subcortical connectivity maps, three views, MNI coordinates. The seed is highlighted with green contours. **List of abbreviations:** Brainstem nuclei used as seeds (marked with a red bracket): Superior Colliculus (SC), Inferior Colliculus (IC), Ventral Tegmental Area-Parabrachial Pigmented Nucleus (VTA-PBP), Microcellular Tegmental Nucleus-Prabigeminal nucleus (MiTg-PBG), Lateral Parabrachial Nucleus (LPB), Medial Parabrachial Nucleus (MPB), Vestibular nuclei complex (Ve), Parvicellular Reticular nucleus Alpha (PCRtA), Superior Olivary Complex (SOC), Superior Medullary Reticular formation (sMRt), Viscero-Sensory Motor nuclei complex (VSM), Inferior Medullary Reticular formation (iMRt), Raphe Magnus (RMg), Raphe Obscurus (ROb) and Raphe Pallidus (RPa). Additional brainstem nuclei used as targets: Median Raphe nucleus (MnR), Periaqueductal Gray (PAG), Substantia Nigra-subregion1 (SN1), Substantia Nigra-subregion2 (SN2), Red nucleus-subregion1 (RN1), Red Nucleus-subregion2 (RN2), Mesencephalic Reticular formation (mRt), Cuneiform (CnF), Pedunculotegmental nuclei (PTg), Isthmic Reticular formation (isRt), Laterodorsal Tegmental Nucleus-Central Gray of the rhombencephalon (LDTg_CGPn), Pontine Reticular Nucleus, Oral Part-Pontine Reticular Nucleus Caudal Part (PnO-PnC), Locus Coeruleus (LC), Subcoeruleus nucleus (SubC), Inferior Olivary Nucleus (ION), Caudal-rostral Linear Raphe (CLi-RLi), Dorsal raphe (DR), and Paramedian Raphe nucleus (PMnR).

In Figure 11A we compared the connectivity results obtained at 7 Tesla and 3 Tesla. We remark a good similarity, with an R index of 0.80 for the comparison between the two brainstem-to-brain connectome matrices. The index decreased to 0.77 when computed on seeds-to-brainstem or seeds-to-others subnetwork matrices. In addition, as shown in Figure 11B, the percentage of common links between 7 Tesla and 3 Tesla data decreased by increasing the statistical threshold; for a Bonferroni-corrected threshold p<0.05, 89.4% of connections were in common between the two data-sets for the seeds-to-brain matrix. This value was equal to 72% for seeds-to-brainstem links and to 93% for seeds-to-others links.

**Figure 11.**
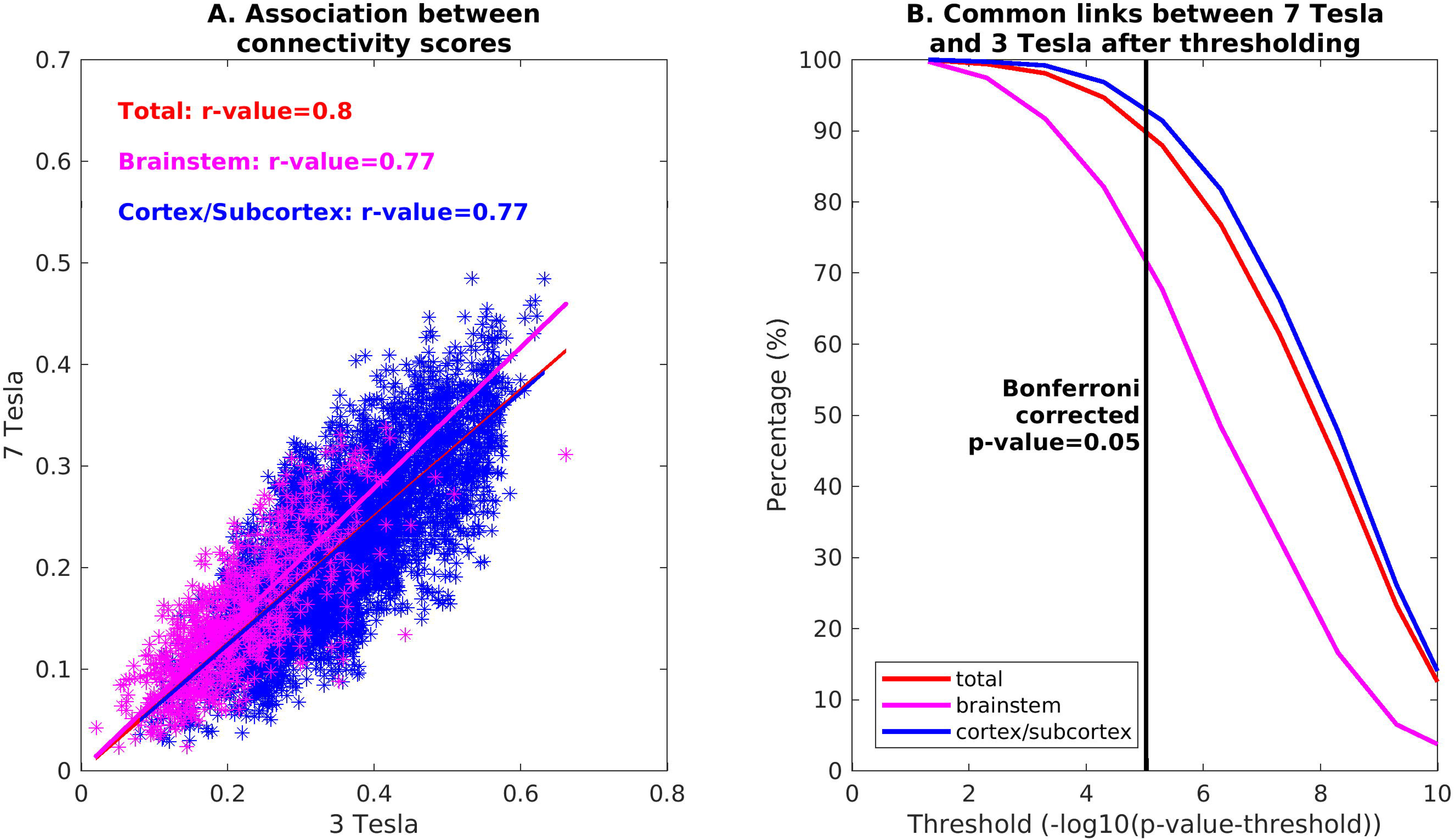
Comparison between 3 Tesla and 7 Tesla connectivity results: **A)** Association between connectivity scores (averaged across subjects and unthresholded) across scanners for the full seeds-to-brain connectome (black), seeds-to-brainstem connectome (magenta), and seeds-to-others (blue). **B)** Percentage of links in common between 7 Tesla and 3 Tesla results, computed for the full seeds-to-brain matrix (red), seeds-to-brainstem matrix (magenta) and seeds-to-others matrix (blue), with respect to different uncorrected statistical thresholds. The vertical line marks the p=0.05 threshold, Bonferroni corrected.

In Figure 12, we show the diagrams depicting significant connections within the limbic/autonomic/nociceptive network (A), the vestibular network (B), and the vestibular nuclei projections to autonomic regions (C). Both networks displayed dense intraconnectivity, and the vestibular nuclei in the brainstem showed low-to-medium interactions with all the autonomic regions except for RMg.

**Figure 12.**
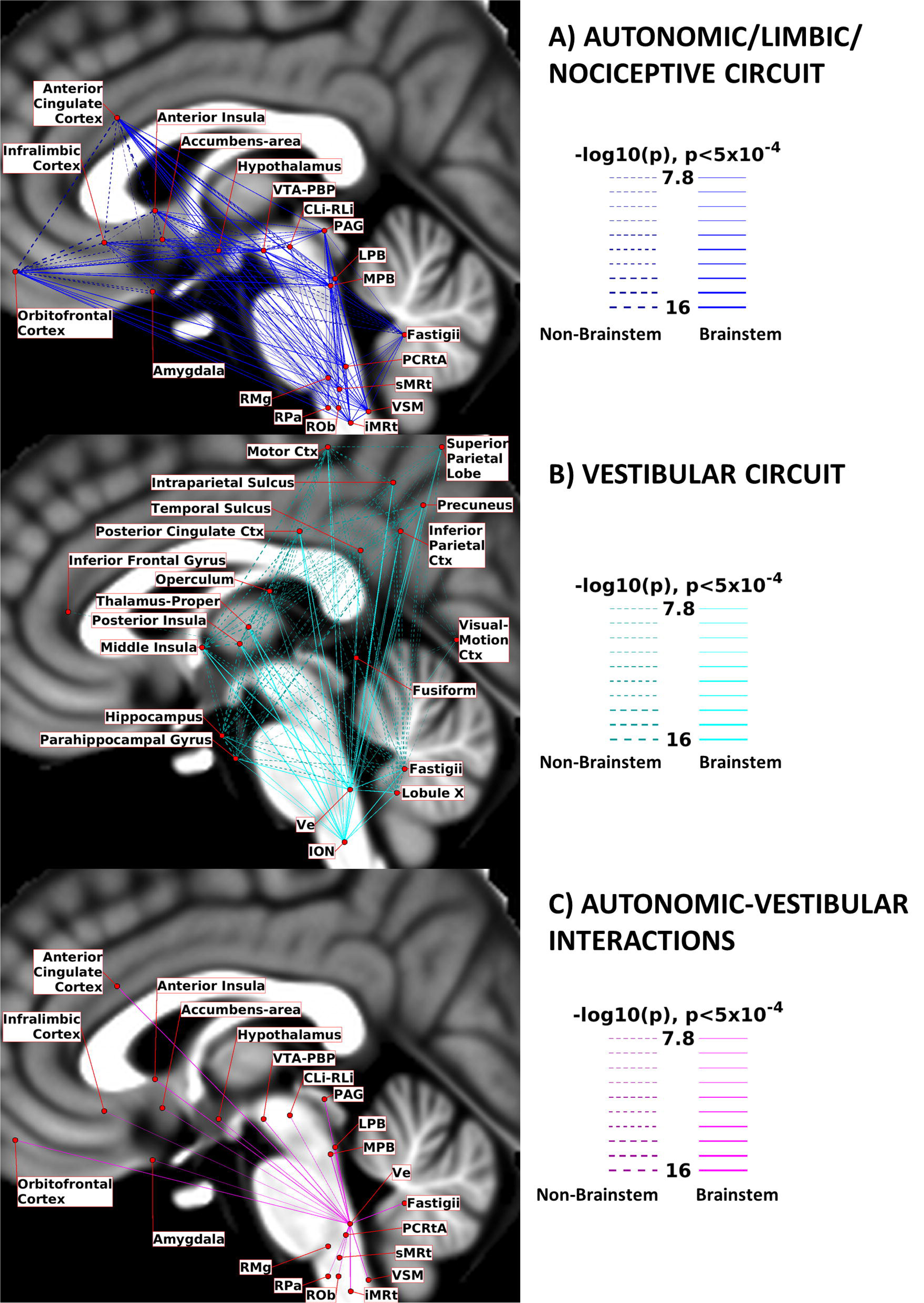
Circuit Diagrams: we summarize here the links among nodes of interest for A) the Autonomic/Limbic/Nociceptive circuit, B) the Vestibular circuit and C) the interactions between Ve and autonomic nodes. Brainstem-to-brainstem connections are displayed with solid lines, while other connections are displayed with dashed lines. Line thickness is proportional to its –log10(p) value, thresholded at p<0.0005, Bonferroni corrected. The nodes positions (region centroids) were projected on the sagittal slice at coordinate x=-2 of the T_1_-weighted MNI template, with small adjustments in their positions to avoid node/link crowding.

## Discussion

### Summary of Results

In this work, we used 7 Tesla functional MRI of 20 healthy subjects to delineate an in-vivo functional connectome of 15 brainstem nuclei involved in central autonomic function and in sensory processing. We refined the coregistrations between functional and structural images to millimetric precision. We used the Pearson’s linear correlation coefficient to outline functional connectivity between our set of seeds, delineated from multi-contrast 7 Tesla MRI [Bianciardi et al., 2015; García-Gomar et al., 2019; Singh et al., 2020b], and a set of targets derived from a complete cortical and subcortical parcellation [Destrieux et al., 2010; Pauli et al., 2018]. We presented the connectome for each of the 15 left or midline seeds, along with a voxel-based linear regression analysis of the whole brain. We unfolded dense brainstem-to-brain links with remarkably low inter-subject variability and high degree of (mirrored) symmetry between left and right seeds. Moreover, by providing function-based circuit diagrams, we observed dense interconnectivity among regions belonging to the same functional circuit. Finally, we investigated the translatability of our study to conventional fMRI acquired at 3 Tesla, with favorable performance.

### Comparison of Connectome Links with Existing Literature

In the following, we discuss the fMRI-based connectomes of arousal and sensory brainstem nuclei in light of available literature. Specifically, we discuss the connectomes displayed in Figures 3-10 (i.e. for bilateral seeds, connectomes of left seeds). Since the laterality index was shown to be low for all nuclei, similar results were obtained for right seeds.

The **Superior Colliculus (SC)** is part of the brainstem sensory network. Its main role is to integrate multisensory information to orient the head and the body towards objects of interest., It is also heavily interconnected with the arousal-motor network [Olszewski and Baxter, 2014]. We observed functional connectivity with CnF and mRt, in line with the mainly-ipsilateral descending pathway targeting emotional-motor nuclei CnF and mRt to trigger head movements in response to aversive stimuli [May, 2003]. Specifically, the pathway with mRt is crucial for the generation of horizontal saccades [Wang et al., 2013]. Strong bilateral connectivity is shown for the thalamus, which is reported in literature to act as a relay in its mediodorsal and intralaminar nuclei for SC to send information mainly towards primary and secondary visual cortices and the medial temporal cortex [Olszewski and Baxter, 2014]. Very sparse functional connections were achieved for occipital regions, still we can observe significant ipsilateral connectivity to the occipital pole (primary visual area). Other relevant links in our connectome of SC were those bilateral with LDTg and PnO, PAG and midbrain raphe nuclei CLi-RLi and DR, which further locate the SC within the sensory network interconnected with the arousal-motor network. Connectivity to amygdala [McFadyen et al., 2019] was confirmed in our functional data; yet, we could not observe functional connectivity to SN (the GABAergic inhibitory projections from SN to SC act to suppress inappropriate saccades [May, 2003]). Literature in primates also reports descending projections to SC from midtemporal areas, motor and premotor cortices [May, 2006], which in our connectome were present along with connectivity to insular and cingulate regions.

The **Inferior Colliculus (IC)** is another crucial node of the brainstem sensory network, which integrates auditory information and participates in the generation of the auditory motor reflex. First, we observe the strong bilateral connectivity to the thalamus, in line with known ipsilateral and crossed projections to its medial geniculate bodies. From there, auditory information is expected to travel to primary and secondary auditory cortices and other association cortices [Olszewski and Baxter, 2014]: in the context of a dense connectivity with parietal, temporal and frontal lobes, we confirm the presence of projections to the superior temporal gyrus for the primary auditory cortex, and report other strong links with precuneus, orbitofrontal, parahippocampal, insular and cingulate regions. As confirmed in our connectome, IC displayed functional connectivity with motor areas such as SN and pallidum, confirming a role for the external nucleus of IC not only in auditory processing but also in initiating the motor response [Casseday et al., 2002]. Its role within a sensory-motor circuit was further confirmed by the presence of functional connectivity to arousal-motor areas such as CnF, mRt, LDTg-CGPn and PnO-PnC. In our IC connectome, we could not identify the expected connection with SOC [Link and Sloan, 2003].

The Ventral Tegmental Area (VTA) is a set of nuclei involved in motivation, reward and arousal, exerting mainly-dopaminergic inputs to several brain regions deputed to decision-making. The Parabrachial Pigmented Nucleus (PBP) is its biggest nucleus, entirely dopaminergic [Olszewski and Baxter, 2014]. Within the brainstem, we observed strong functional connectivity of the **Ventral Tegmental Area - Parabrachial Pigmented Nucleus (VTA-PBP)** with PAG and DR, both responsible for glutamatergic and GABAergic inputs to VTA neurons. The same role is played by PTg and LDTg, which displayed strong ipsilateral functional connectivity, yet weaker contralateral connectivity [Morales and Margolis, 2017]. In the frontal cortex, we obtained the strongest connections with the anterior cingulate cortex and the pericallosal sulcus, in line with literature specifying bidirectional links with the medial prefrontal cortex [Lavin, 2005]. Within subcortical nuclei, the VTA-PBP displayed strong bilateral connectivity with thalamus, putamen, pallidum and cerebellum. Pallidum is known to project to VTA, along with hypothalamus and ipsilateral accumbens [Morales and Margolis, 2017], which all showed high functional connectivity with VTA-PBP. Other strong functional connectivity values of VTA-PBP to arousal/sleep-motor brainstem nuclei such as mRt, PnO, SubC, iMRt indicates a possible role for VTA-PBP in the initiation of arousal/sleep-motor responses. We also report a very strong ipsilateral connectivity to the brainstem motor nucleus RN1. Interestingly, as shown in Figure S6 in supplementary materials, VTA-PBP was a network hub with the highest score in Betweenness Centrality: this indicates its role in sharing information with different networks, connecting otherwise disconnected brain nodes. This is confirmed by the fact that VTA-PBP was the brainstem nucleus showing in Figure 12A the strongest connections to cortical regions, apparently working as a relay center between the brainstem autonomic/limbic/nociceptive network and the cortical limbic regions deputed to decision making, reward and higher-order limbic processing.

The Microcellular Tegmental Nucleus is a small structure that lies between mRt and the Parabigeminal nucleus, a brainstem visual center which is thought to modulate visual attention. In this study these two regions are merged in the **Microcellular Tegmental Nucleus – Parabigeminal Nucleus (MiTg-PBG)** region. The MiTg was introduced in the human brainstem atlas by [Paxinos et al., 2012] but it is still understudied in the literature. For PBG, our connectome did not display the expected connection with SC, while we observed connectivity to amygdala, which should mediate rapid responses to visual threats [Olszewski and Baxter, 2014]. In this context, it is easy to understand the observed functional connectivity of MiTg-PBG with limbic (PAG, parabrachial nuclei) arousal (DR, mRt, CnF, PnO-PnC) and autonomic-motor nuclei (iMRt, sMRt, VSM), some of which have been reported in cat studies [Baleydier and Magnin, 1979].

The **Lateral Parabrachial Nucleus (LPB)** operates as an interface between the viscero-sensory nuclei of the medulla and autonomic/limbic forebrain structures [Olszewski and Baxter, 2014]. Inputs from viscero-sensory regions such as the caudal solitary nucleus and nucleus ambiguus were indeed confirmed in our connectome by significant functional connectivity with VSM (including the Solitary nucleus), iMRt and sMRt (containing nucleus ambiguus). Furthermore, we observed functional connectivity of LPB with autonomic/limbic forebrain structures such as the hypothalamus and nucleus accumbens (i.e. basal forebrain). These pathways are reported in the literature to be reciprocal and topographically arranged [Olszewski and Baxter, 2014], and have been confirmed in both rat and human studies [Fulwiler and Saper, 1984; de Lacalle and Saper, 2000]. Moreover, we observed strong functional connectivity of LPB with the thalamus, in line with rat studies demonstrating visceral and nociceptive inputs from the LPB to the thalamus, further relayed to autonomic cortical areas such as the insular cortex [Olszewski and Baxter, 2014]. Significant links with arousal nuclei of the brainstem, namely LDTg-CGPn, PnO-PnC, CnF, mRt, DR and CLi-RLi, are in line with the role of LPB as a promoter of arousal responses to stimuli via projections to orexin nuclei in the hypothalamus [Arima et al., 2019]. In addition, strong connections to ipsilateral Ve is at the basis of limbic-vestibular interactions defined in animal studies [Balaban, 2004].

Similar to the LPB, the role of the **Medial Parabrachial Nucleus (MPB)** is at the interface between medullary and forebrain regions; yet it receives prevalently gustatory stimuli, instead of viscero-sensory ones, and projects mainly to the limbic forebrain [Olszewski and Baxter, 2014]. In contrast to LPB, MPB is reported to directly project to cortical regions without having thalamus as relay: in our results we observe both “direct” functional connectivity with the anterior insular cortex, cingulate and infralimbic cortex, and strong connectivity with the thalamus, maybe mediated by descending cortico-thalamic pathways. Expected connectivity with amygdala was present but weak, while the functional connectivity to LC was strong [Yang et al., 2021]. The connectivity to medullary reticular formation nuclei (iMRt and sMRt) and VSM was bilateral, and in line with the presence of afferents from the rostral gustatory division of the solitary nucleus (included in VSM) [Olszewski and Baxter, 2014]. Finally, we also observed functional connectivity of the MPB with iMRt and sMRt, in line with the evidence that pharynx, larynx and esophagus are controlled by the MPB via projections to the nucleus ambiguus (comprised within iMRt and sMRt).

The **Vestibular Nuclei Complex (Ve)** is the center of a broad vestibular network encompassing several brain regions [Singh et al., 2021b]. The ION is reported to play a role as a relay between Ve and the cerebellum [Olszewski and Baxter, 2014], where lobule X and fastigial nuclei are part of the vestibular network. As described in [Indovina et al., 2020], this network extends to several cortical and subcortical regions, i.e. the posterior cingulate cortex (in particular the visual area in the cingulate sulcus as also reported in studies on macaques [De Castro et al., 2021] and humans [Smith et al., 2017], posterior and middle insular cortex, inferior frontal gyrus, parahippocampal gyrus, fusiform gyrus, visual-motion cortex, primary motor cortex, inferior parietal cortex, precuneus, intraparietal sulcus, operculum, thalamus and hippocampus. As visible in the vestibular circuit diagram (Figure 12B), Ve displayed functional connectivity with all these regions, with particularly high connectivity values with the thalamus, parietal operculum and posterior insular cortex, mid-posterior cingulate cortex and cerebellum, as also visible from the Ve circular connectome (Figure 6, top panel). The inferior frontal gyrus was the only node disconnected from Ve, and from almost all the rest of the network. Occipital and temporal regions displayed lower interconnectivity within the network, while the functional connectivity of the motor cortex was very strong, with the thalamus being a possible mediator due to its strong connectivity both with this cortical region and with brainstem nodes. Beyond the connectivity within the vestibular network, we also investigated the autonomic/limbic-vestibular interactions, because adverse vestibular-autonomic/limbic interactions [Brandt, 1996; Bronstein, 2004; Indovina et al., 2015; Jacob et al., 2009; Staab, 2012; Staab et al., 2014; Staab et al., 2017] appear to precipitate and perpetuate chronic vestibular disorders, crucially underlying the pathophysiologic process of these disorders. In Figure 12C we observed strong interactions with autonomic/limbic/nociceptive brainstem nuclei, the strongest being crucially with iMRt and parabrachial nuclei and the weakest with raphe nuclei (RPa, ROb, CLi-RLi). The strongest functional connectivity with autonomic/limbic/nociceptive cortical nodes was with the orbitofrontal and anterior cingulate cortex and fastigial nuclei. Mild connectivity is present with the amygdala (the latter was also mildly connected to MPB and LPB). As shown in Figure S6 in supplementary materials, Ve displayed the lowest Clustering Coefficient and the highest Participant Coefficient among seeds, which is an indication of the high interconnectivity with distant cortical regions in different modules.

The **Superior Olivary Complex (SOC)** is the first relay center of the auditory pathway, directly integrating information from cochlear nuclei in the pons to the IC. Notably, in the context of a quite sparse functional connectome of SOC, we observed a contralateral projection to IC. The SOC also includes the human equivalent of the trapezoid body, which has a primary role in the spatial localization of sounds [Paxinos et al., 2012]. The bilateral connection of SOC to limbic and motor nuclei iMRt and MPB could reflect its participation in the initiation of the auditory reflex and conscious perception.

The **Superior Medullary Reticular formation (sMRt)** is a medullary region which includes [Garcia Gomar et al., 2021] the compact part and the superior portion of the semicompact part of the nucleus ambiguus, the gigantocellular reticular nucleus, the parvicellular reticular nucleus, the intermediate reticular nucleus, the dorsal paragigantocellular nucleus and the facial motor nucleus [Paxinos et al., 2012]. The connectome of the ambiguus nucleus is scarcely described in literature; yet the functional connectivity of sMRt with VSM is in line with the key role of ambiguus nucleus in the control of cardiac activity [Saper and Stornetta, 2015]; our connectome analysis, also confirmed the functional connectivity of sMRt with Ve, which might underlay autonomic-vestibular interactions [Balaban and Beryozkin, 1994]. The gigantocellular nucleus is a node of the somatic-motor system associated with the generation of motor patterns for jaw, head, tongue and face, as well of neuromodulation [Olszewski and Baxter, 2014]. Links to motor nuclei such as VSM are thus expected, as well as to limbic/neuromodulatory areas such as PAG and to multisensory areas such as SC [Holstege, 1991]. Interestingly, in our connectome, we reported functional connectivity with PAG and VSM, but this was not the case for SC. We also observed connectivity with other nuclei of the emotional-motor network such as CnF and MPB.

The loose part and the inferior portion of the semicompact part of the ambiguus nucleus, as well as the retroambiguus nucleus, the ventral and dorsal medullary reticular nucleus, the intermediate reticular nucleus, noradrenaline cells A1 and adrenaline cells C1 are included instead in the **Inferior Medullary Reticular formation (iMRt)** [Garcia Gomar et al., 2021]. The retroambiguus nucleus interacts with the ambiguous nucleus for the control of swallowing and speech. Functionally, the retroambiguus nucleus contains phasic respiratory modulated neurons and plays thus a specific role in respiration control [Jones et al., 2012]. Anatomically, it appears as a column of cells caudal to the loose part of nucleus ambiguus [Watson, 2012]; its circuitry is still understudied in literature, with cat studies reporting interactions of the caudal part with PAG, amygdala and hypothalamus [Holstege, 1991]. In our connectome, we observed functional connectivity of iMRt with all these regions. The dorsal medullary reticular nucleus is involved in the modulation of spinal nociceptive processing. In our results, we observed functional connectivity with ION, VSM, PAG, SN1 and SN2 that were reported in studies on rats [Leite-Almeida et al., 2006], as well as links to sMRt in line with projections to the facial motor nucleus and paragigantocellular nucleus. In the diencephalon, we found very strong connections with the thalamus and the cerebellar cortex.

The **Viscero-Sensory Motor Nuclei Complex (VSM)** includes the solitary nucleus, the dorsal motor nucleus of the vagus and the hypoglossal nucleus [Singh et al., 2021b]. The hypoglossal nucleus is mainly known for its moto-neurons controlling tongue muscles; interestingly, we observed functional connectivity with iMRt, which includes the nucleus retroambiguus and the dorsal medullary reticular nucleus, involved in swallowing, respiration and vocalization. Further, the observed functional connectivity of VSM with parabrachial nuclei is in line with the interaction of the hypoglossal nucleus with the Kölliker-Fuse for the integration of respiratory signals [Olszewski and Baxter, 2014], while the functional connectivity to VTA could be explained by their common input from the cortical motor tongue area [Paxinos et al., 2012]. The solitary nucleus is the primary brainstem sensory integration area for taste, gastrointestinal, baroreceptor, respiratory and cardiovascular information from peripheral receptors, and as such it is expected to target several autonomic and premotor nuclei. We observed functional connectivity of VSM to medullary reticular formation nuclei (iMRt, sMRt) that are crucial for vital reflexes, such as vomiting, and for cardiovascular and respiratory responses [Olszewski and Baxter, 2014]. Moreover, we observed significant functional connectivity between VSM and parabrachial nuclei, which confirms the role of the latter as relay centers between the solitary nucleus and autonomic forebrain structures [Fulwiler and Saper, 1984; de Lacalle and Saper, 2000; Olszewski and Baxter, 2014], as well as collect peripheral information to guide respiratory activity. Catecholaminergic links to PAG were also confirmed in the VSM connectome [Saper and Stornetta, 2015]. the dorsal motor nucleus of the vagus is a visceral efferent strongly interconnected with the central autonomic network including hypothalamus, insular cortex, amygdala, PAG and Parabrachial nuclei [Olszewski and Baxter, 2014]. All these connections were visible in our results.

The **Raphe Magnus (RMg)** is a mainly serotonergic nucleus with a major role in the modulation of nociception and CO_2_ chemoreception [Saper and Stornetta, 2015]. In this study, RMg showed very sparse functional connectivity; we could not confirm connectivity with other brainstem nuclei except for slightly-suprathreshold connectivity with left iMRt and right LDTg, which might be related respectively to the serotonergic descending pathway and to modulatory interactions with the limbic system. The observed functional connectivity with the Dorsal Medullary Reticular Nucleus, which is included in iMRt, was reported in cats [Leite-Almeida et al., 2006]. Further, we observed functional connectivity with the PAG, which is reported to excite RMg and LC to block pain transmission with its anti-nociceptor neurons [Michael-Titus et al., 2010]. With regard to the cortex, we confirmed rat studies reporting connections with the insular cortex, while we did not obtained links with the perirhinal cortex [Hermann et al., 1997].

The **Raphe Pallidus (RPa)** is the smallest raphe nucleus and, as Raphe Obscurus, it plays a modulatory role on brainstem motor nuclei. Structural connections for the RPa have been studied in rats: connectivity with PAG, and LPB (including the Kölliker-Fuse) were confirmed in the functional connectome of RPa, while connectivity with sMRt (which includes the lateral paragigantocellular reticular nucleus) was not significant [Hermann et al., 1997]. We also observed a link with iMRt (which includes part of the nucleus ambiguus). Further, we observed links of RPa to the hypothalamus, which participates in thermoregulation, and bilateral connectivity with the amygdala, but none with the limbic-motor nucleus SubC [Olszewski and Baxter, 2014]. The observed functional connectivity to DR and CLi-RLi suggests a modulatory role of RPa in arousal as well. Links to Ve were bilateral, indicating its possible role in modulating vestibular processing [Halberstadt and Balaban, 2003].

The **Raphe Obscurus (ROb)** provides modulatory serotonergic input to brainstem motor nuclei such as VSM, nucleus accumbens and pre-Boetzinger [Saper and Stornetta, 2015]. We observed indeed bilateral connections to iMRt and sMRt, as well as a contralateral link to VSM. Literature specific to ROb is quite sparse, yet we confirmed a bilateral link with Ve [Halberstadt and Balaban, 2003] and PAG [Olszewski and Baxter, 2014]. We also found functional connectivity of ROb with LDTg, PnO and LPB, consistent with the involvement of ROb in modulating the actuation of emotional and autonomic processes.

The **Alpha Part of the Parvicellular Reticular Nucleus (PCRtA)** [Garcia Gomar et al., 2021] is a relatively understudied part of the Parvicellular Reticular Nucleus (PCRt), introduced in [Paxinos et al., 2012]. PCRt (included in sMRt in this study) has been described as a multipurpose region: it integrates sensory information from cranial nerves and projects it to oral-motor regions that control blinking, salivation, mastication, and similar movements; it also participates in cardiovascular and respiratory regulation [Shammah-Lagnado et al., 1992]. Interestingly, in line with the functions of PCRt, in our connectome, we observed functional connectivity of PCRtA with motor regions (cerebellum, basal ganglia, VSM, iMRt, ION, mRt, PnO-PnC) and with arousal/autonomic regions (VSM, iMRt, VTA-PBP, mRt, LDTg-CGPn and PnO-PnC).

### Circuit Diagrams

In Figure 12A, the autonomic/limbic/nociceptive circuit is summarized in a schematic diagram, considering as brainstem autonomic/limbic/nociceptive nodes PAG, VTA-PBP, LPB, MPB, PCRtA, sMRt, iMRt, VSM, RMg, ROb, RPa and CLi-RLi. We note a high degree of interconnectivity among the nodes of this network. Intracortical connectivity was particularly strong among orbitofrontal cortex, anterior cingulate cortex, infralimbic cortex and anterior insula. In Figure 12B, the vestibular circuit is depicted in a schematic diagram, while in Figure 12C we depicted the connections between the vestibular nuclei complex and the autonomic/limbic/nociceptive nodes used in Figure 12A. Regarding these vestibulo-autonomic interactions, we observed functional connectivity of Ve with almost all (except for RMg) autonomic/limbic/nociceptive brainstem nuclei.

### Relevance

The definition of this functional connectome might provide a baseline for future studies investigating alterations in autonomic and sensory function related to pathological conditions. Several studies have already defined functional connections that can be used to build predictive models in the clinical field, for example in studies of schizophrenia, ADHD, major depression, autism, epilepsy, prenatal cocaine exposure and multiple sclerosis [Castellanos et al., 2013].

In addition, being able to describe the functional connectome of multiple brainstem regions provides new insight into the description of segregation and integration among brainstem circuits that operate in a coordinated fashion. For instance, this is the case for the autonomic and vestibular circuits and their interaction, which we further discuss as an example. For decades, neurophysiologic research on recovery from acute vestibular deficits has focused on compensatory mechanisms in the brainstem vestibular nuclei and associated brainstem/cerebellar pathways [Dutia, 2010; Kitahara et al., 1997; Ris et al., 1997]. In contrast, recent prospective clinical studies [Best et al., 2009; Cousins et al., 2014; E.J. Mahoney et al., 2013; Godemann et al., 2005; Heinrichs et al., 2007] identified elevated autonomic arousal and anxiety as the primary predictors of failed recovery and prolonged vestibular symptoms (e.g. dizziness, imbalance, hypersensitivity to motion stimuli) in chronic vestibular disorders following acute vestibular events. Thus, methods able to map in living humans subcortical pathways linking vestibular nuclei to anxiety regions (e.g., amygdala) via parabrachial autonomic nuclei (LPB, MPB) and other associated autonomic nuclei (e.g. PAG, RMg, Solitary Nucleus) are needed. They can expand our knowledge of successful compensation for acute vestibular events versus development of chronic vestibular disorders. Treatment is much less effective after the disorder is established, so a better understanding of its neural mechanisms, offered by advanced imaging methods, could guide development of more effective, even preventative, interventions to reduce its prevalence and disability.

Finally, this functional connectivity study serves as a further validation of the probabilistic MRI template of autonomic and sensory brainstem nuclei [Bianciardi et al., 2015; García-Gomar et al., 2019; Singh et al., 2020b] defined based on multi-contrast 7 Tesla MRI in living humans. It does so by demonstrating the presence of functional connectivity of arousal and sensory brainstem seeds with subcortical and cortical regions expected based on previous animal studies and on the functional circuit definition.

### Novelty

To our knowledge, several resting state functional connections reported in this study were observed for the first time in living humans. This is because studies of the resting state functional connectivity of brainstem nuclei in healthy subjects are sparse. For example, connections of SC with CnF, mRt, LDTg, PnO, PAG, CLi-RLi and DR, as well as connections of IC with SN, pallidum, CnF, mRt, LDTg-CGPn, PnO-PnC are original, given the lack of literature on their functional connectome at rest with other brainstem nuclei. The same applies to the connectivity of SOC with IC, iMRt and MPB; to the connectivity of MiTg-PBG with amygdala, PAG, parabrachial nuclei, DR, CnF, mRt, PnO-PnC, iMRt, sMRt and VSM; to the connectivity of PCRtA with VSM, iMRt, ION, mRt, PnO-PnC, VSM, iMRt, VTA-PBP, mRt, LDTg-CGPn and PnO-PnC; to the connectome of reticular formation nuclei iMRt and sMRT, the former with PAG, amygdala, hypothalamus, ION, VSM, PAG and substantia nigra, the latter with VSM, Ve, PAG; to the connectivity of VSM with iMRt, parabrachial nuclei, VTA, medullary reticular formation nuclei and PAG. VTA-PBP connectivity with PAG, DR, PTg, LDTg-CGPn, mRt, PnO, SubC, iMRt are novel as well, while other subcortical ones are already documented. The connectome of parabrachial nuclei has been investigated only with task-based fMRI [Sklerov et al., 2019], thus, to our knowledge, our findings on their connectivity to the rest of the brain are original as well. For Ve, available literature of the vestibular network only reports the functional connectivity to the whole brainstem as a single node [Noohi et al., 2020] or to roughly divided brainstem macroregions [Kirsch et al., 2016]; thus, we consider our findings of Ve connectivity as original. The functional connectome of RMg has been investigated in [Bianciardi et al., 2016 and Bär et al., 2016], which also report its connections to PAG and insula within the salience network. To these findings we reported novel (slightly-suprathreshold) connectivity of RMg with iMRT and LDTg. Functional connectomes of the other medullary raphe nuclei, RPa and Rob, also represent a novelty.

### Strengths

The strength of this study is rooted on the ad-hoc tools at our disposal. First, we used an ultra-high field MRI scanner equipped with a custom receive coil-array with enhanced sensitivity for subcortical areas, including the brainstem. This allowed us to obtain images with high spatial resolution, yet with whole brain coverage and a reasonable temporal resolution, also thanks to the fast simultaneous-multi-slice acquisition sequence employed. Second, we used a detailed probabilistic MRI-based template of the brainstem comprising 58 nuclei in stereotactic space: thanks to this template we relied on a precise localization of these nuclei, mostly understudied in studies on functional connectivity due to their difficult localization. Third, our experimental protocol was designed to provide a high number of data points (i.e. degrees of freedom), which combined with a good sample size (n = 20 subjects) allowed us to achieve good statistical significance. Fourth, we achieved high accuracy in the coregistration of functional images to stereotactic (MNI) space, which was crucial considering the small size of our brainstem seeds; further, we did not smooth the data to extract the seed average signal time-courses to decrease partial volume effects. Finally, we limited the number of spatial resampling steps (which cause spatial smoothing) by concatenating all the coregistration steps: motion correction, distortion correction, affine registration and nonlinear warp to subject’s structural image, registration of the structural image to the custom template, and registration of the custom template to the MNI standard template.

### Sources of resting state fMRI signal fluctuations

There are several drivers of resting state activity underlying the derived connectomes and diagrams. These include gut-brain, lung-brain, heart-brain, and sensory-organs-brain (e.g. eye-brain) interactions, which involve brainstem nuclei and other brain regions. For example, the impact of oscillations in gastrointestinal activity on the fMRI signal have been investigated via gastric electrical stimulation [Cao et al., 2019]. The latter study highlights the crucial role of the vagus nerve in the modulation of the gut-brain axis, and reports activations in the midcingulate and insular cortex, somatosensory and motor cortices, as well as visual and auditory ones. Interestingly, the connectome of the VSM, a region of interest including the dorsal motor nucleus of the vagus, presented several strong connections with the cortex (cingulate and insular cortex, pericallosal sulcus), thalamus, cerebellum, basal ganglia, as well as with brainstem nuclei, such as the VTA-PBP, IC and PnO-PnC. In addition, gastrointestinal disease has an impact on the connectivity of the default mode network with other brain regions [Skrobisz et al., 2020]. Ascending viscera-sensory pathways through the vagal nerve enter the solitary nucleus (part of VSM) and are further relayed to the parabrachial nuclei, raphe nuclei, locus coeruleus and to the forebrain, which crucially are present in our connectome.

Further, regarding lung-brain interactions, the respiratory-related sensorimotor activity in medullary and pontine respiratory groups is expected to modulate the signal correlations within the autonomic/limbic/nociceptive circuit. The well-note network of respiratory rhythm pattern generators is then integrated by several chemosensitive brainstem nuclei which provide additional feedback, e.g., LC, medullary raphes, pre-Boetzinger complex and solitary nucleus, the latter relaying information also from peripheral chemoreceptors and pulmonary stretch receptors [Nattie and Li, 2012]. The solitary nucleus (part of VSM) is a relay station for the baroreflex and for the whole heart-brain axis as well: concurrent recording of muscle sympathetic neural activity and fMRI revealed coupled activity at rest in the solitary nucleus, ventrolateral medulla, PAG, insula and hypothalamus [Macefield and Henderson, 2019]. In the medulla, connectivity of VSM with sMRt reflects afferents of solitary nucleus to vagal cardiomotor neurons in the nucleus ambiguus. Functional connectivity within the network deputed to control of cardiac activity was found to be nonlinearly related to heart rate [de la Cruz et al., 2019]. Fluctuations in heart rate and respiration at rest are expected to be an important driver of the observed correlations among these regions, described in our autonomic/limbic/nociceptive circuit.

Moreover, gazing and spontaneous saccades [Koba et al., 2021] have an impact on the functional connectome, specifically on motor and sensory-motor networks, as well as the dorsal attention network, While unfortunately these studies did not investigate brainstem regions, the trajectories of saccades with eyes closed are different than with eyes open [Becker and Fuchs, 1969] and a different effect on the functional connectome may be expected. Interestingly, in the connectome of SC, strong connections with mRt and CnF were visible. mRt has a key role in the generation of saccades, and CnF is a sensory-motor nucleus. Rather, connections with auditory regions, such as the IC, were not confirmed.

### Limitations

In spite of optimized coregistration and distortion correction methods, we expect the presence of residual distortions, small mis-alignment and signal dropout in certain areas, artifacts that are particularly strong in ultra-high field MRI. All subjects showed adequate alignment between the brainstem edges of the functional image and those of the standard template, in particular for the medulla, midbrain, and dorsal pons, while some residual distortions could be spotted on the rostral part of the pons for six out of 20 subjects. Luckily, the 15 seeds of interest were not located in the anterior pons. Nonetheless, connectivity to amygdala and ventral orbitofrontal targets might be affected by the dropout present in fMRI in these regions [Ojemann et al., 1997].

In the current work, to compute the association among time-courses, we employed Pearson correlation, which has at least three drawbacks. First, it is a measure of linear correlation, which is unable to capture nonlinear mixing of inputs at a node that results in output signals. For instance at some nodes, nonlinear mixing of descending multimodal inputs is implemented by crucial subcortical and brainstem nodes, such as the amygdala [Gothard, 2020] or the superior colliculus [Kasap and van Opstal, 2017], to encode affective and social behaviours or fast and simple motor output. These nonlinear dynamics are expected at the level of spike rate transformations: their impact on the BOLD signal is a nontrivial issue though, because of its complex and still unclear relationship with local field potentials, action potentials, and glutamate release [Vanni et al., 2015]. Nevertheless, propagation of these small-scale nonlinear dynamics onto larger scale BOLD dynamics are not expected to be captured by linear correlation measures between BOLD signal changes in functionally-connected regions [DiNuzzo et al., 2019]. Second, Pearson correlation is affected by indirect (polysynaptic) connectivity, which limits the interpretation of resting state networks. For example, the observed fMRI connectivity between VSM and accumbens might be an indirect connection, resulting from oscillations in gastrointestinal and cardiovascular activity, which modulate both the direct expected connectivity between VSM and parabrachial nuclei, as well as the direct connectivity between parabrachial nuclei and acccumbens [Fulwiler and Saper, 1984; de Lacalle and Saper, 2000; Olszewski and Baxter, 2014]. A third drawback is that Pearson correlation restricts the estimated connectivity to instantaneous effects. However, in literature we can find evidence of several sources of correlation that expand to non-zero time lags and may exhibit dynamic changes in time. Future fMRI connectivity analyses based on measures of nonlinear relation [Reshef et al., 2011], multivariate conditional mutual information [Sundaram et al., 2020], measures of partial coherence [Wang et al., 2016], dynamic approaches to connectivity [Hutchison et al., 2013], as well as nonparametric tests of statistical significance that avoid underlying distributional assumptions of the data might improve the accuracy and specificity of connectivity values.

Our human connectomes were rich of expected connectivity pathways based on animal studies, yet they also lacked some relevant connections: this was the case for the amygdala, which showed modest connectivity within both the autonomic network and the vestibular network. For example, the MPB displayed high connectivity to the LC, cingulate and insular cortices, but weak connectivity with the amygdala. In general, the amygdala is a difficult area to be studied with MRI, because it is a small region affected by high spatial distortions and signal dropout, due to its proximity to the paranasal sinuses. Moreover, its signal fluctuations are also confounded by neighbouring large veins draining distant brain regions [Boubela et al., 2015], and by respiratory-related magnetic field changes due to chest motion, which are more prominent in deep brain regions than in the cortex. Further work with distortion-reduced fMRI methods [Wang et al., 2019] and higher order shimming [Stockmann and Wald, 2018] might improve the connectivity profile of the amygdala.

In our study, we asked our participants to keep their eyes closed during fMRI without falling asleep, and we checked their awake condition at the beginning of each run. The choice of acquiring resting state data on subjects with their eyes closed (or open) is nontrivial, and is known to affect the connectivity results [Agcaoglu et al., 2019; Bianciardi et al., 2009; Costumero et al., 2020; Patriat et al., 2013]. For example, the connectivity of the auditory and sensorimotor networks are increased during the eyes open condition compared to the eyes closed one [Agcaoglu et al., 2019; Patriat et al., 2013]; yet, literature results vary for the connectivity of occipital areas, some reporting increased connectivity during the eyes closed condition [Bianciardi et al., 2009; Costumero et al., 2020], and others during the eyes open condition [Agcaoglu et al., 2019; Patriat et al., 2013]. Our connectomes show in general lower connectivity of seeds to occipital regions. Despite that, there is no agreement in literature, the use of the eyes closed condition is still widespread. We recognize that the generalizability of connectivity results is limited by the adopted condition, and by considerations emerging from these studies.

In this work we used minimum (for the chosen number of slices and spatial resolution) TRs of 2.5 s (for 7 Tesla fMRI) and 3.5 s (for 3 Tesla MRI), as well as a band-pass filter with cut-off frequencies of 0.01 Hz and 0.1 Hz. This is a standard approach for conventional fMRI connectivity analyses focused on evaluating signal fluctuations occurring over five seconds to tens of seconds. Moreover, the employed cut-off frequencies have been broadly used to clean the signal from non-neuronal causes and improve the temporal signal-to-noise-ratio [Lee et al., 2013]. For example, using a lowpass filtering cutoff of 0.1 Hz allows to use a TR up to 5 s (which is the TR that corresponds to the 0.1 Hz Nyquist frequency). For these reasons, in the literature there are also 3 Tesla fMRI studies with longer TRs, for example of 6 seconds [Horovitz et al., 2008]. The use of long TRs has some drawbacks, e.g. the chance of aliasing cardiac artefacts on the same frequencies of the BOLD signal [Huotari et al., 2019]. Nevertheless, algorithms for physiological noise correction, such as RETROICOR, model these artefacts including their aliasing effects. Lately, there is increased interest in the use of shorter TR, mainly enabled by the use of simultaneous-multi-slice imaging methods and of low spatial resolutions, to investigate faster BOLD signal fluctuations and dynamic changes in functional connectivity in health (e.g. saccade generation) and pathological conditions, such as epilepsy or central apnea [Sahib et al., 2018].

Our preprocessing pipeline made use both of temporal filtering and of nuisance regression. In our work, this was particularly efficient thanks to the concurrent measure of physiological signals, i.e. of cardiac and respiratory related signal fluctuation, and to the availability of a large number of data points. The correct preprocessing procedure for connectivity analysis is still debated in literature, and this debate focuses mainly on motion correction, temporal filtering and nuisance regression [Bright and Murphy, 2015; Gargouri et al., 2018]. Head motion has a great impact on the BOLD signal and various solutions have been proposed, mainly related to the regression of additional timecourses originated from the derivative and energy of the 6 motion timeseries (translation and rotation for each of the three axes) [Maknojia et al., 2019; Parkes et al., 2018], which was implemented in the current work. Other sources of debate are the nature and amount of nuisance regressors that should be used in order to remove non-neural BOLD fluctuations, such as physiological noise. In particular, average timecourses extracted from macroregions or the global signal have been long used as nuisance regressors, but these standard procedures have been lately questioned [Caballero-Gaudes and Reynolds, 2017], along with the standard filtering applied to resting state fMRI [Davey et al., 2013]. Despite this, no consensus has been reached; we provided in the method section a detailed description of the employed preprocessing step, thus a highly reproducible pipeline, as suggested in [Lindquist, 2020].

### Conclusions

In conclusion, we described in detail the functional connectome of 15 autonomic and sensory brainstem nuclei in living humans, reporting general considerations about the whole connectivity matrix, as well as discussing specific connections on the basis of existing literature. We confirmed the high interconnectivity among nuclei belonging to different (autonomic, vestibular) networks, and the feasibility of extracting nuclei-specific information through ultra-high field resting-state fMRI and the use of a probabilistic brainstem nuclei atlas. In addition, we showed results suggesting a good translatability for these results in conventional (e.g. 3 Tesla) scanners, especially for brainstem-cortical functional connectivity and to a lesser extent for brainstem-brainstem functional connectivity. This functional connectome might serve as a baseline for human studies of pathological conditions involving autonomic, limbic, nociceptive and sensory functions.

## Supporting information

https://figshare.com/s/43422a6b1e2f0edbe4d1

## Acknowledgements

This work was funded by grants from the National Institutes of Health (NIBIB-K01EB019474, NIDCD-R21DC015888, NIA-R01AG063982), from the MGH Claflin Award, Harvard Mind Brain Behavior, and by the Italian Ministry of Health PE-2013-02355372

## List of Abbreviations

PTg: Pedunculopontine Tegmental Nucleus
LDTg-CGPn: Laterodorsal Tegmental Nucleus and Central Gray of the Rhombencephalon
PnO-PnC: Pontine Reticular Nucleus, Oral Part and Pontine Reticular Nucleus, Caudal Part
LC: Locus Coeruleus
SubC: Subcoeruleus
ION: Inferior Olivary Complex
CLi-RLi: Caudal Linear Raphe and Rostral Linear Raphe
SC: Superior Colliculus
IC: Inferior Colliculus
VTA-PBP: Ventral Tegmental Area and Parabrachial Pigmented Nucleus
MiTg-PBG: Microcellular Tegmental Nucleus
LPB: Lateral Parabrachial
MPB: Medial Parabrachial
Ve: Vestibular nucleus
SOC: Superior Olivary Complex
sMRt: Superior Medullary Reticular Formation
iMRt: Inferior Medullary Reticular Formation
VSM: Viscerosensory Motor Nucleus
RMg: Raphe Magnus
ROb: Raphe Obscurus
RPa: Raphe Pallidus
PCRtA: Parvicellular Reticular Nucleus, Alpha Part

