## Supplementary material for "Functional connectome of brainstem nuclei involved in autonomic, limbic, pain and sensory processing in living humans from 7 Tesla resting state fMRI": https://figshare.com/s/43422a6b1e2f0edbe4d1

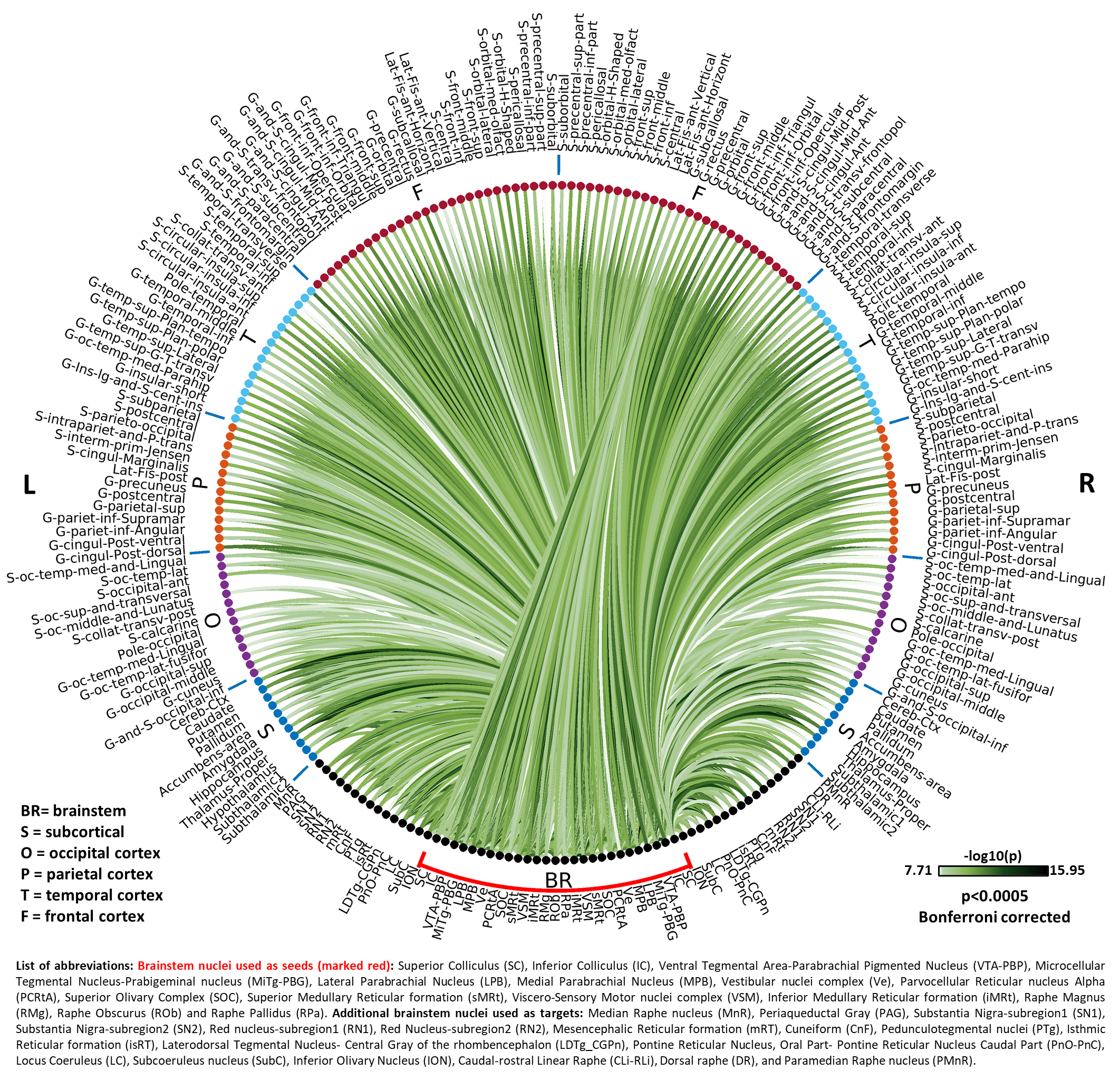

**Figure 1S) Seeds-to-Brain circular connectome:** circular plot displaying the –log_10_(p-value) extracted from the group-level region-based analysis of seeds-to-brain functional connectivity (p<0.0005, Bonferroni corrected).

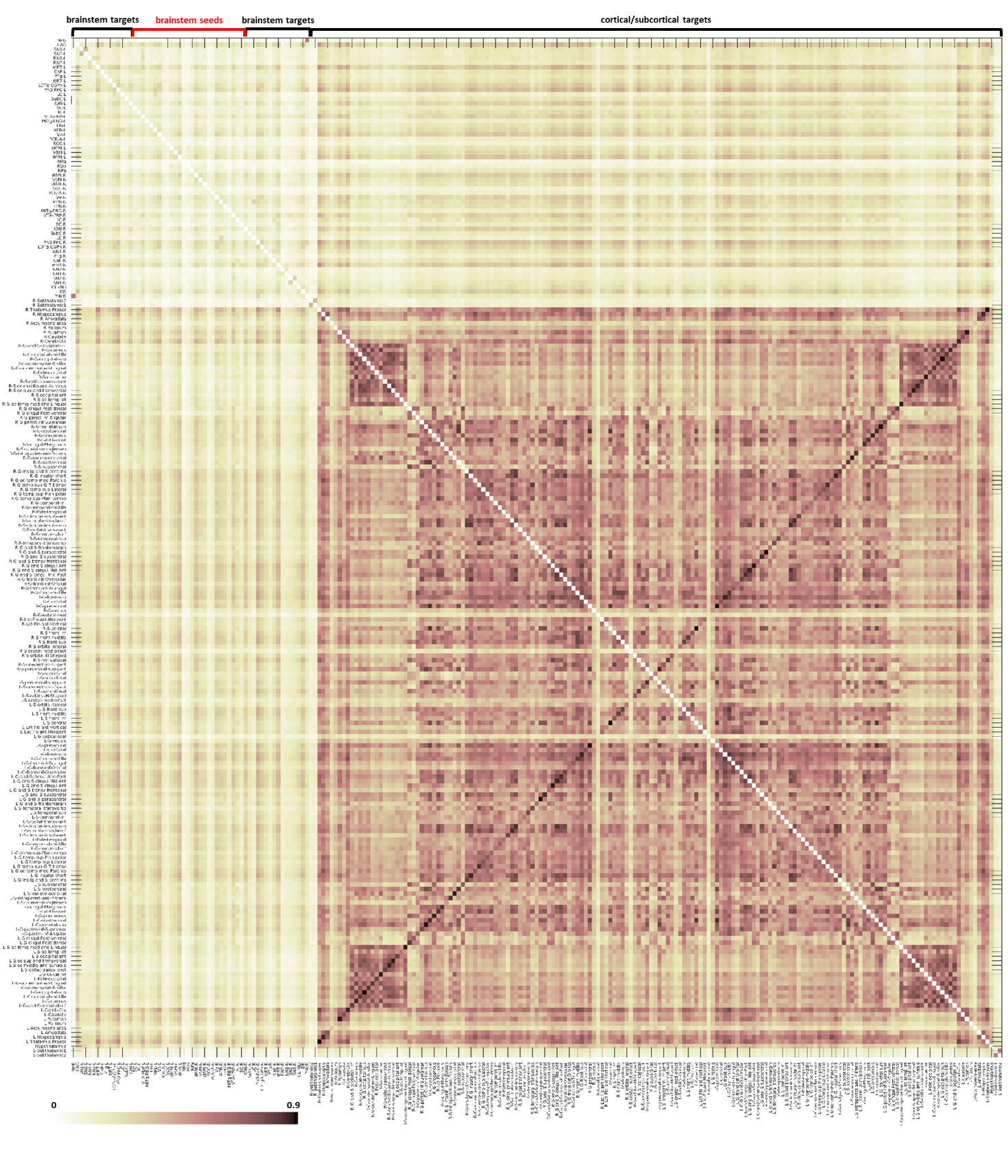

**Figure S2) Brain-to-Brain Correlation Coefficient matrix, average values across subjects**

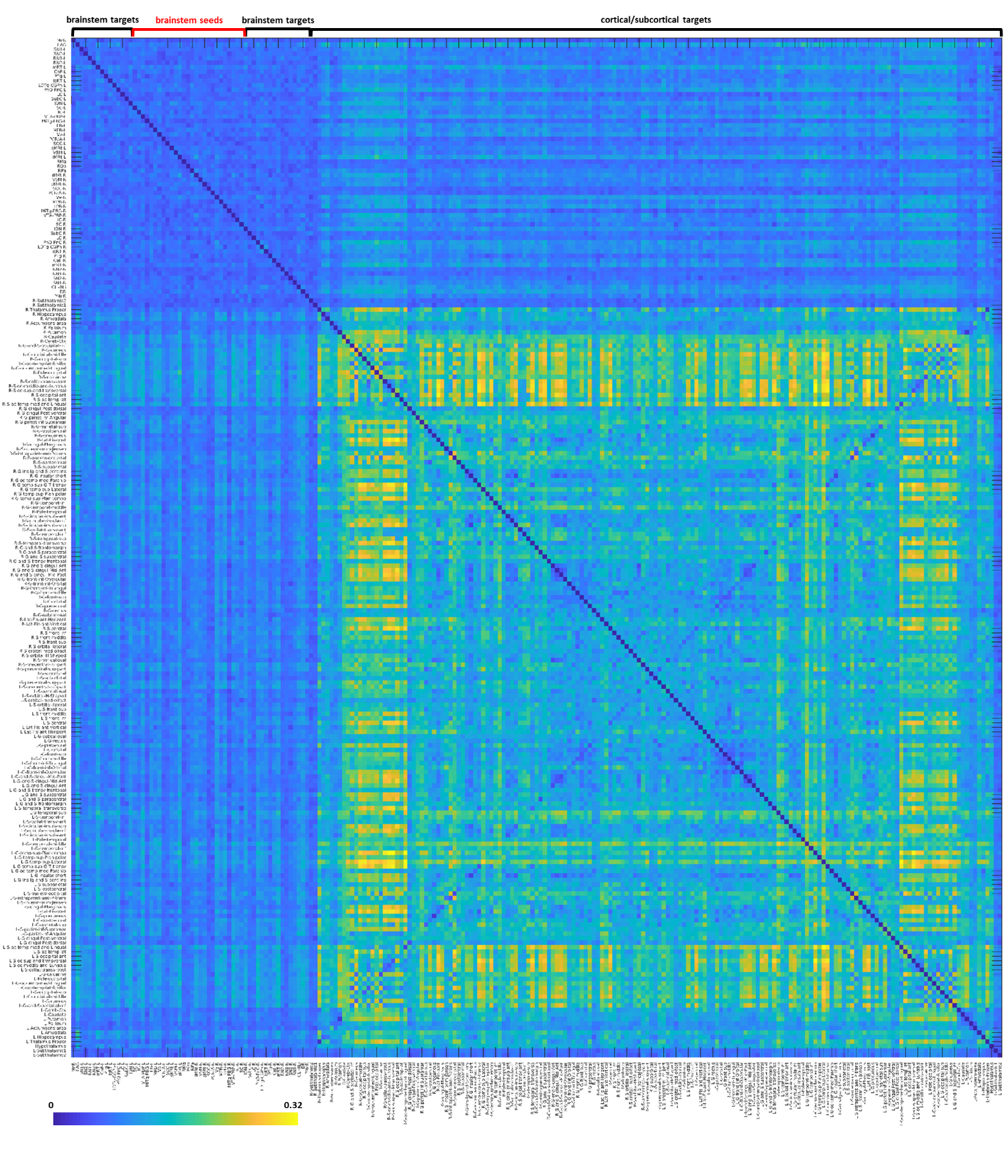

**Figure S3) Brain-to-Brain Correlation Coefficient matrix, standard deviation across subjects**

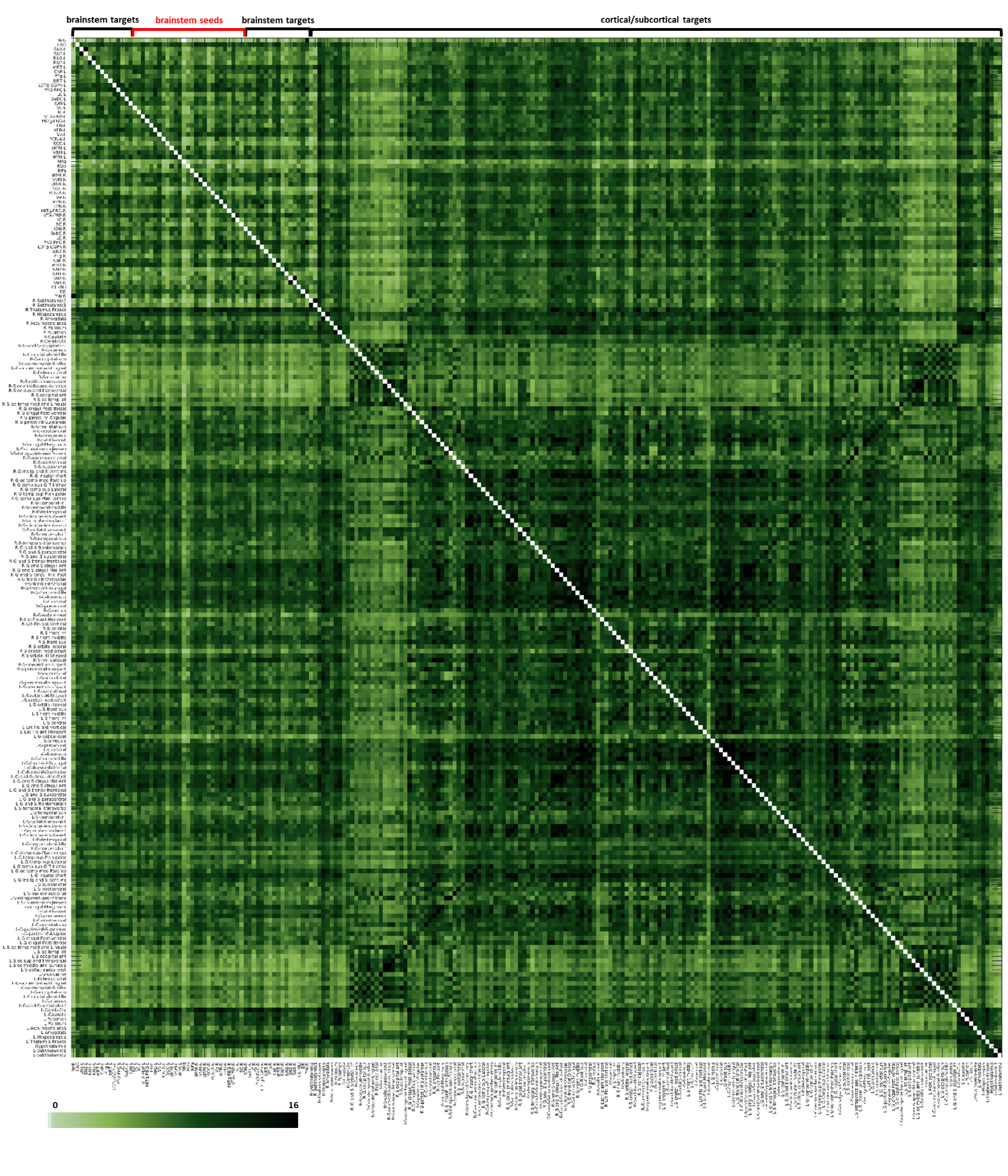

**Figure S4) Brain-to-Brain Correlation Coefficient matrix, unthresholded -log10(p-value)**

### Graph analysis

#### Materials and Methods

Graph analysis metrics were computed using the GRETNA (GRaph thEoreTical Network Analysis) Matlab toolbox (<http://www.nitrc.org/projects/gretna/>, [Wang et al., 2015]). We extracted global graph measures (assortativity, rich club, synchronization, hierarchy, clustering coefficient, characteristic path length, gamma, lambda, sigma, global efficiency and local efficiency) on the whole brain-to-brain network and on the seeds-to-brainstem and seeds-to-cortex subnetworks. Further, for each seed and each target region, we extracted nodal measures (betweenness centrality, degree centrality, local efficiency, clustering coefficient, shortest path length, normalized participant coefficient). Rich club was computed considering the binary Degree Centrality averaged on the whole network or on the subnetwork of interest. To compute the Participant Coefficient, the network was divided into seven communities (seeds, brainstem targets, and then subcortical, occipital, temporal, frontal and parietal cortical targets). Occasionally, nodal measures were compared across subgroups of labels using a two-sample t-test. Finally, for each bilateral seed, we computed (in Matlab) a laterality index defined as the difference between the binarized connectome of the left seed and the mirrored binarized connectome of the right seed, divided by the number of active links, thus scoring 0% for perfectly symmetric connectivity and 100% for perfectly asymmetric one.

#### Results

In Figure S5, global graph measures are displayed for the seeds-to-seeds connectome (panel A) and the cortex-to-cortex connectome (panel B). With respect to the cortex-to-cortex network, the seeds-to-seeds network showed anti-assortative topology (seeds-to-seeds: -0.3, cortex-to-cortex: 0.08), lower global clustering coefficient (seeds-to-seeds: 0.53, cortex-to-cortex: 0.56) and a tendency towards a hierarchical structure (seeds-to-seeds: 0.15, cortex-to-cortex: -0.06). Small-world measures Lambda, Gamma and Sigma were all close to one in both cases. Efficiency was higher in the cortex both at global (seeds-to-seeds: 7.3, cortex-to-cortex: 9.3) and local level (seeds-to-seeds: 8.5, cortex-to-cortex: 10.07). The lowest section of panel C reports the laterality index of bilateral seeds: no seed exceeded 18% laterality, with only SOC, SC, PCRtA and MiTg-PBG exceeding 10%.”). We also moved the section relative to nodal measures (“In Figure S6 we report nodal graph measures for all seeds, averaged across bilateral seeds. These measures were computed for all target regions too (see Supplementary Figures S7-11). For all seeds, Betweenness Centrality was consistently lower than the network average (35.76); the maximum among seeds was reached by VTA-PBP, with a value of 5, then MPB and VSM. Ve, iMRt, VSM, MPB and VTA-PBP had Degree Centrality values higher than the network average, thus were defined as network hubs. RMg, SOC, ROb and SC displayed the highest values for Local Efficiency, while only Ve, iMRt and RPa failed to reach the network average. Clustering coefficient was higher (p<0.001, two-sample t-test) in the brainstem (mean±std 0.562±0.027) with respect to the rest of the brain (0.527±0.033). Shortest Path Length results were similar to those of Local Efficiency. Nodes with the highest Normalized Participant Coefficient were Ve, iMRt, sMRt, MPB, LPB and VSM.

#### Discussion

The graph analysis of the connectivity matrix revealed that the autonomic and sensory functional networks did not show a significatively clusterized topology, and consequently did not show not even for the cortical subnetwork the expected, still debated, small-world organization hypothesized at least for the forebrain [Hilgetag and Goulas, 2016], even if the condition on ≈1 on Lambda is met [Bassett and Bullmore, 2006]. Note that small-world topology has been mostly observed in anatomical networks, while in functional networks results varied depending on the parcellation used and in particular on the number of nodes [Wang et al., 2009]. Interestingly, the brainstem seeds-to-seeds subnetwork showed disassortative mixing (ASS=-0.3), in line with literature on functional cortical connectomes [Lim et al., 2019], while the cortico-cortical subnetwork displayed positive and close to zero assortativity. The Clustering Coefficient was slightly higher in the brainstem than in the cortex, which demonstrated a slightly more hierarchical organization. Based on nodal measures, Ve, iMRt, VSM, MPB and VTA-PBP were defined as network hubs, having Degree Centrality higher than the network average value. Being a brain networks hub is a relevant feature, which can be altered in pathological or perturbed conditions [Vatansever et al., 2020]. In general, the brainstem nodes used as seeds showed lower Betweenness Centrality than the rest of the network: higher values were instead obtained for known relay centers such as the thalamus, hypothalamus and cerebellum, as well as by some orbitofrontal cortical nodes (see Supplementary Materials).

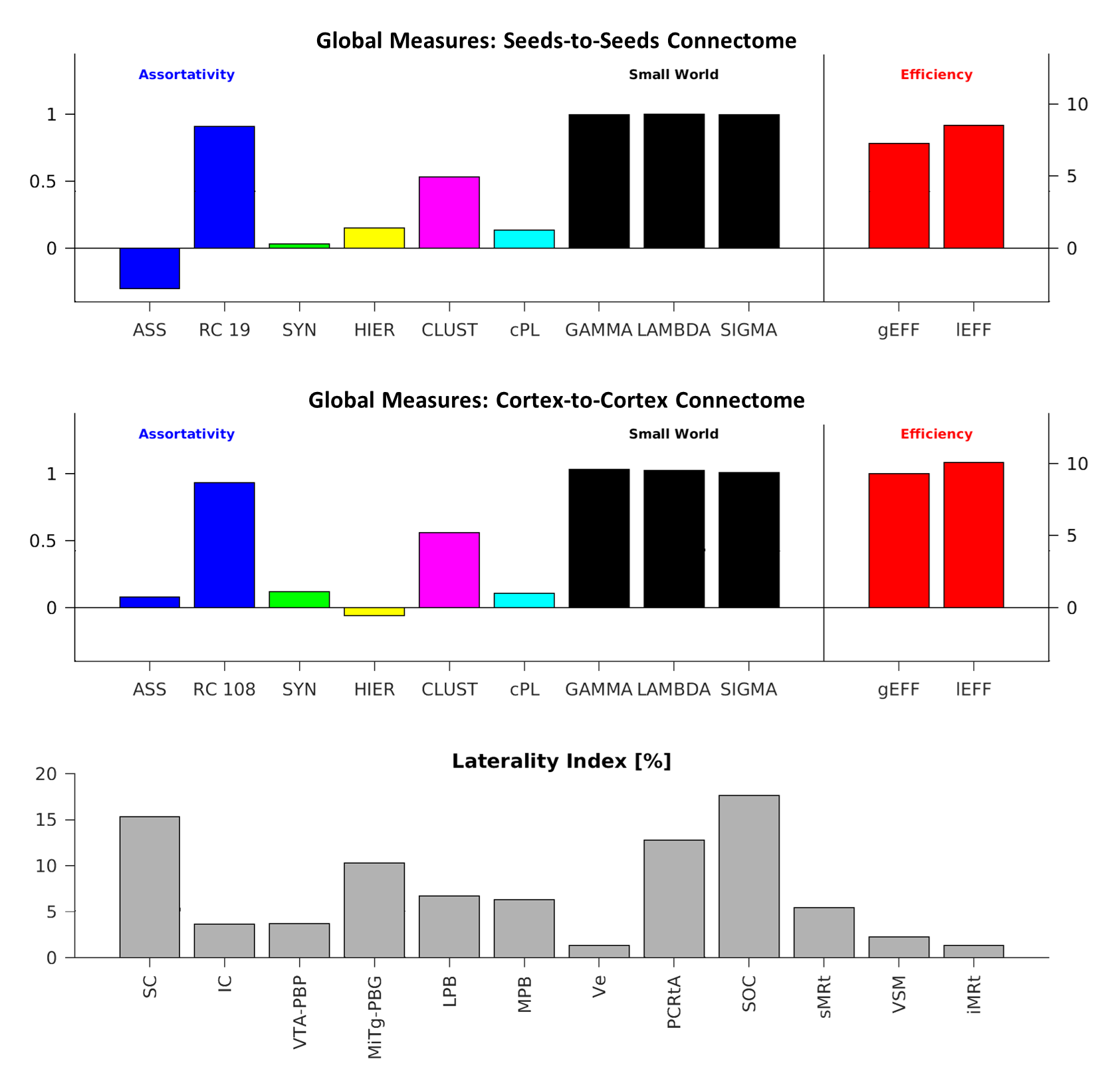

**Figure S5) Global Graph measures and nodal laterality indexes**: (top) global graph measures extracted with GRETNA for (top) seeds-to-seeds connectome and (middle) cortex-to-cortex connectome: Assortativity (ASS); Rich Club (RC); synchronization (SYN); hierarchy (HIER); Characteristic Path Length (cPL); GAMMA; LAMBDA; SIGMA; global Efficiency (gEFF); local Efficiency (lEFF). (Bottom): laterality index for bilateral seeds (0% = perfect symmetry): SC, IC, VTA-PBP, MiTg-PBG, LPB, MPB, Ve, PCRtA, SOC, sMRt, VSM, iMRt.

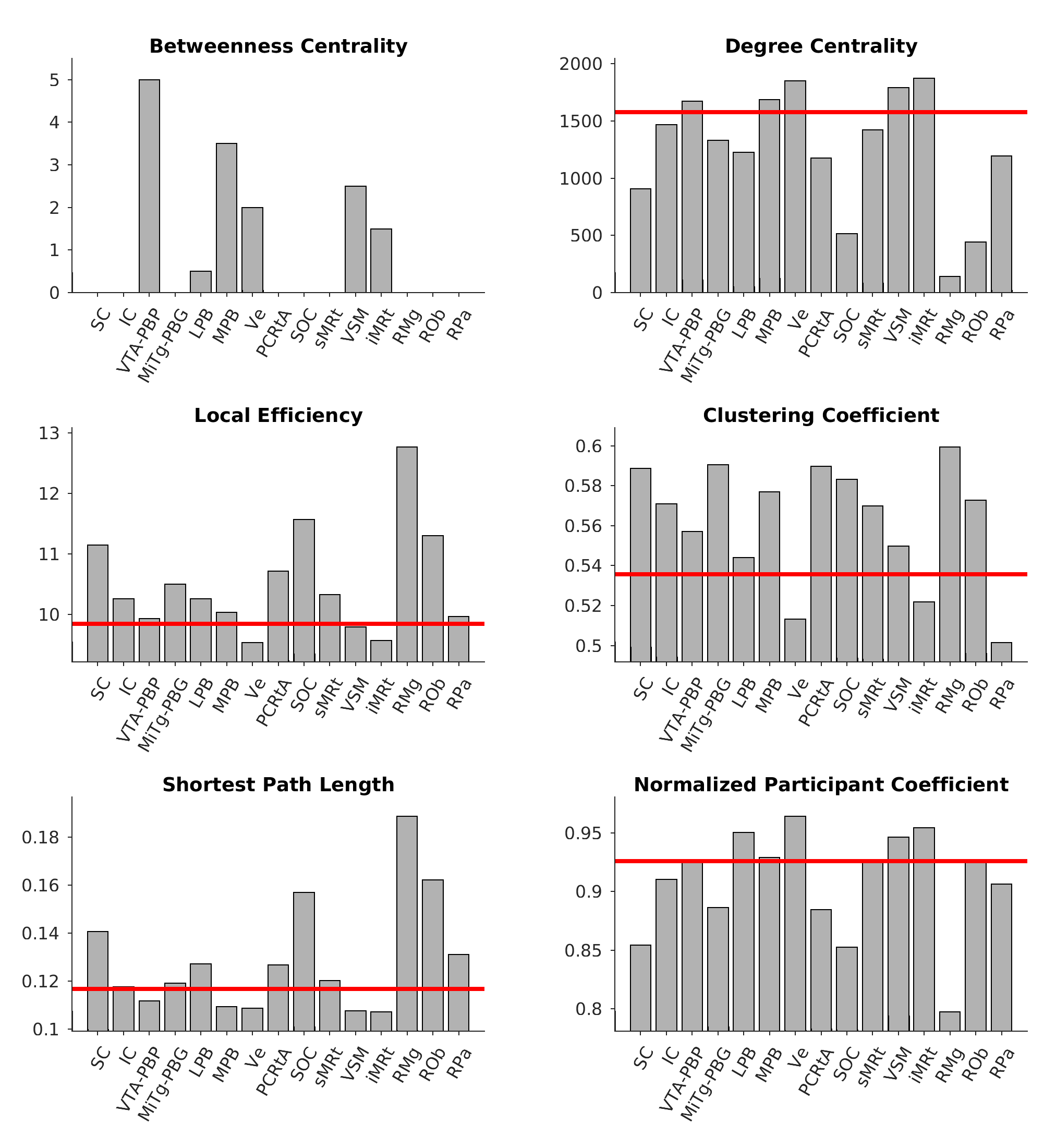

**Figure S6)** **Nodal graph-analysis metrics of the functional connectome seeds**: Red lines mark the whole-network-averaged values. **Betweenness Centrality**: VTA-PBP, MPB and VSM were the nodes that mostly contributed to the efficiency of the network, by promoting short paths. Note that the network average was 35.76, and thus it was not displayed in the bar plot. **Degree Centrality**: network hubs were Ve, iMRt, VSM, MPB and VTA-PBP (i.e. with degree centrality higher than the network average). **Local Efficiency**: RMg, SOC, Rob and SC displayed the highest values of local efficiency. **Clustering Coefficient**: All the seeds, except for Ve, iMRt and RPa, exceeded the network average. **Shortest Path Length**: Results were similar to those of Local Efficiency. The smaller values were found for iMRt, VSM, Ve, MPB and the highest for RMg, ROb, SOC, and SC. **Normalized Participant Coefficient**: interestingly, nodes contributing the most to inter-community communication were Ve, iMRt, sMRt, LPB, MPB, and VSM. **List of abbreviations:** Superior Colliculus (SC), Inferior Colliculus (IC), Ventral Tegmental Area-Parabrachial Pigmented Nucleus (VTA-PBP), Microcellular Tegmental Nucleus-Parabigeminal nucleus (MiTg-PBG), Lateral Parabrachial Nucleus (LPB), Medial Parabrachial Nucleus (MPB), Vestibular nuclei complex (Ve), Parvicellular Reticular nucleus Alpha (PCRtA), Superior Olivary Complex (SOC), Superior Medullary Reticular formation (sMRt), Viscero-Sensory Motor nuclei complex (VSM), Inferior Medullary Reticular formation (iMRt), Raphe Magnus (RMg), Raphe Obscurus (ROb) and Raphe Pallidus (RPa).

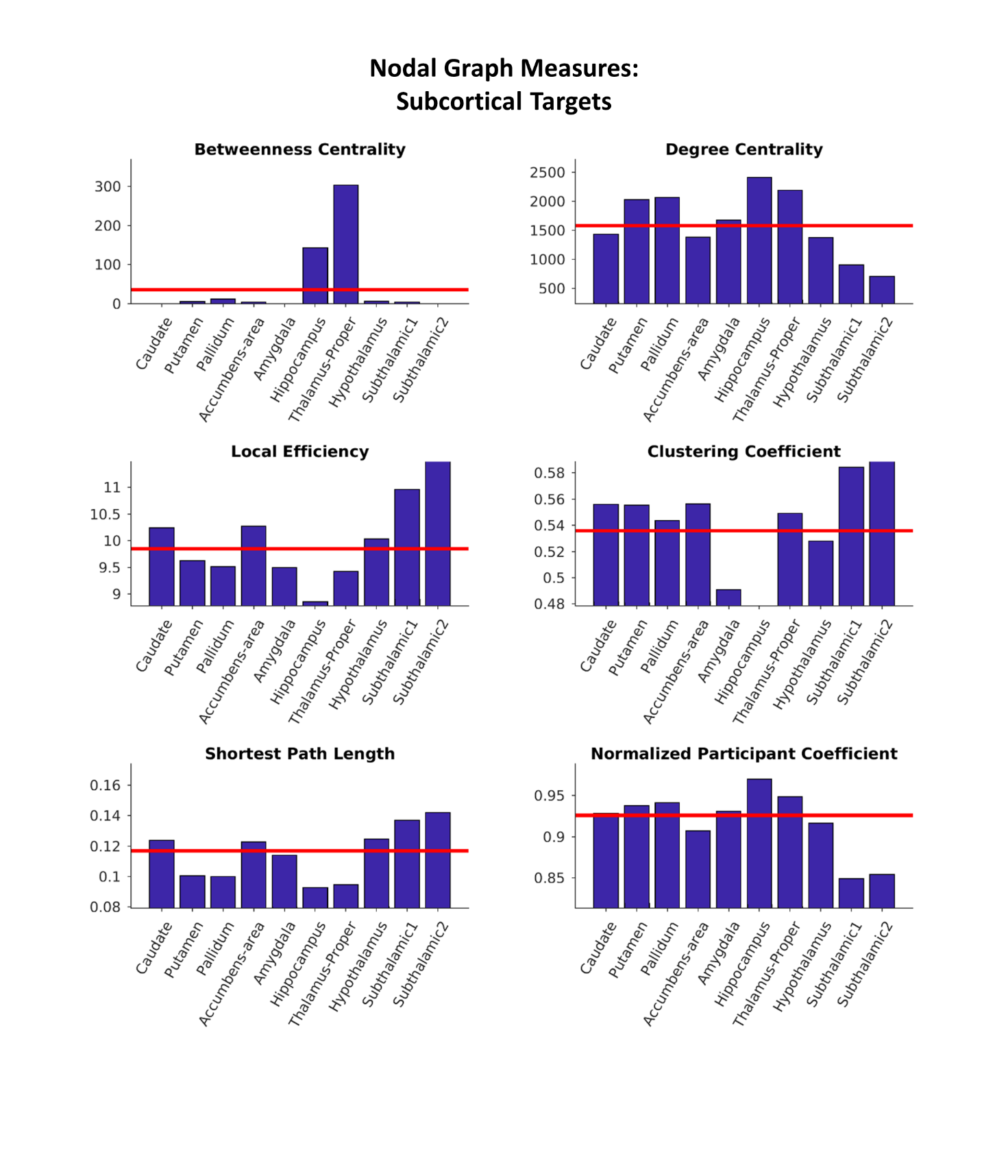

**Figure S7)** **Nodal graph-analysis metrics of the functional connectivity of subcortical targets**. Red lines mark the whole-network-averaged values.

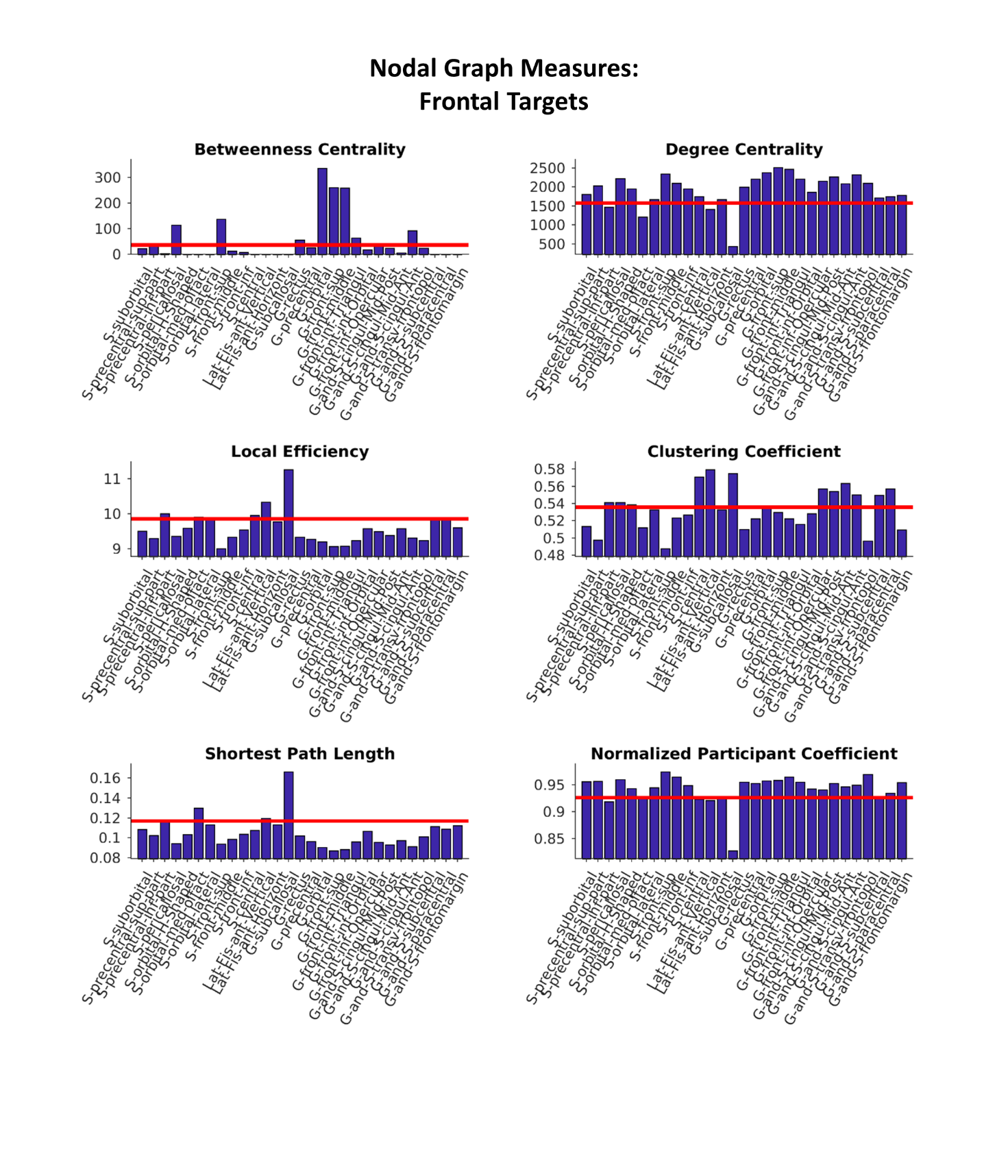

**Figure S8)** **Nodal graph-analysis metrics of the functional connectivity of targets of the frontal lobe**. Red lines mark the whole-network-averaged values.

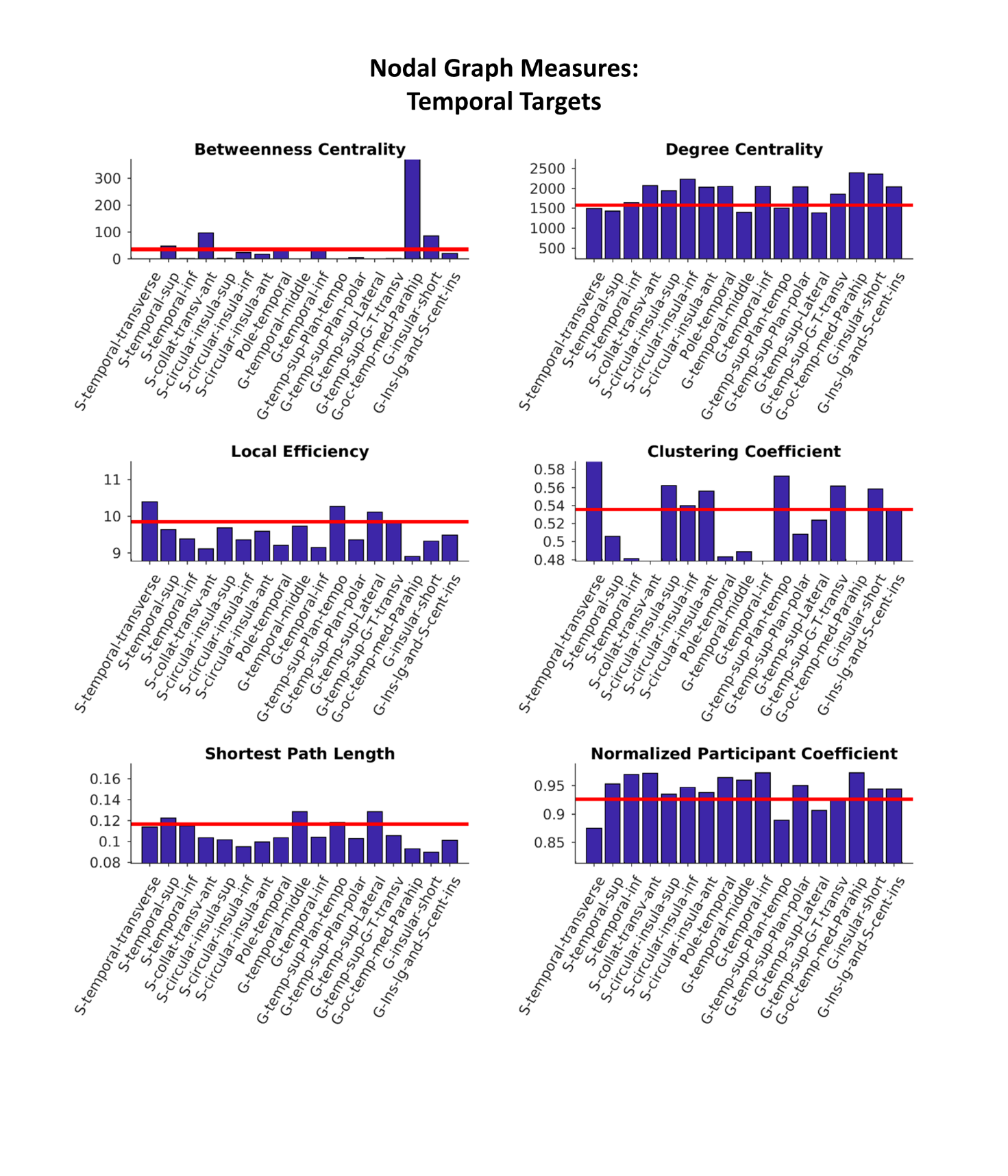

**Figure S9)** **Nodal graph-analysis metrics of the functional connectivity of targets of the temporal lobe**. Red lines mark the whole-network-averaged values.

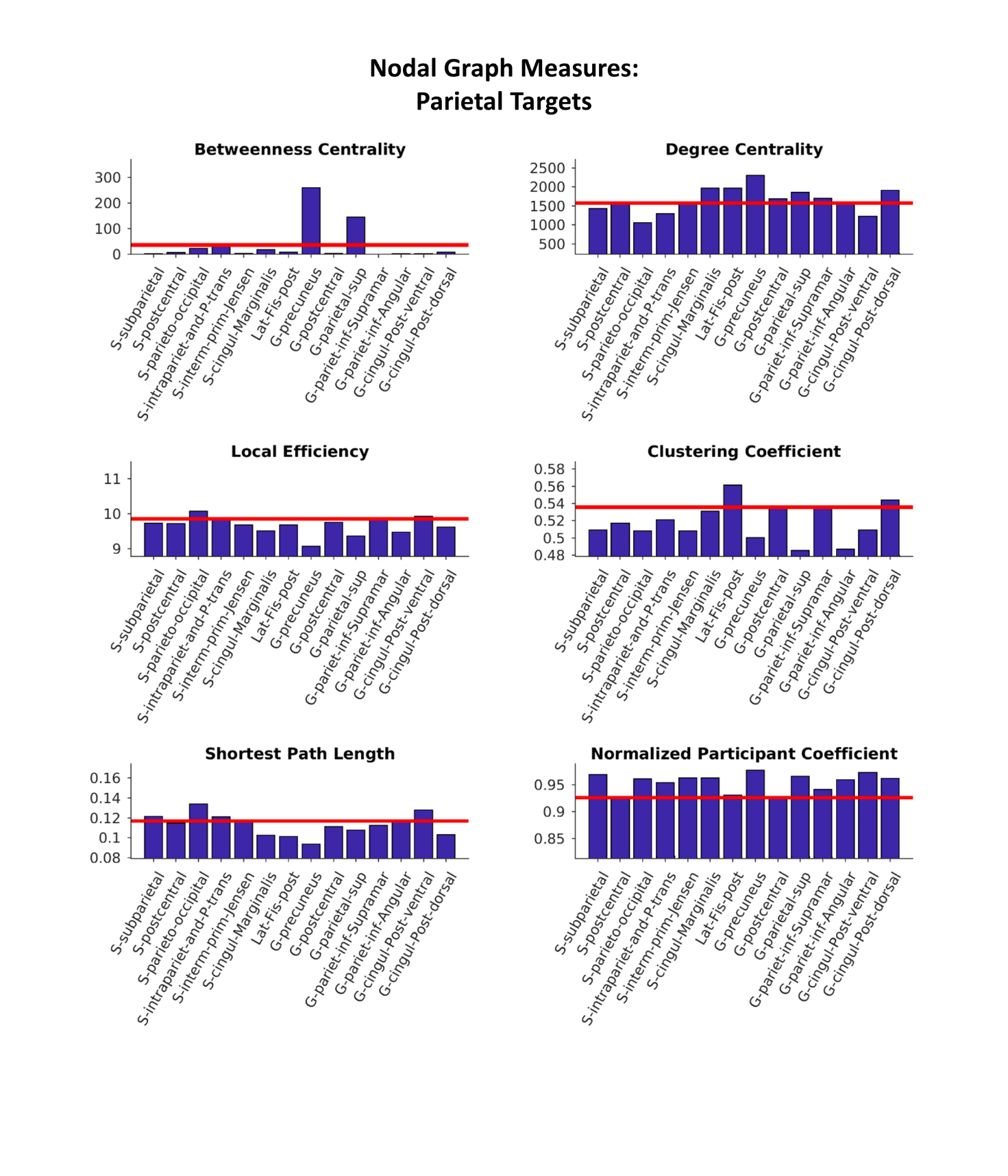

**Figure S10)** **Nodal graph-analysis metrics of the functional connectivity of targets of the parietal lobe**. Red lines mark the whole-network-averaged values.

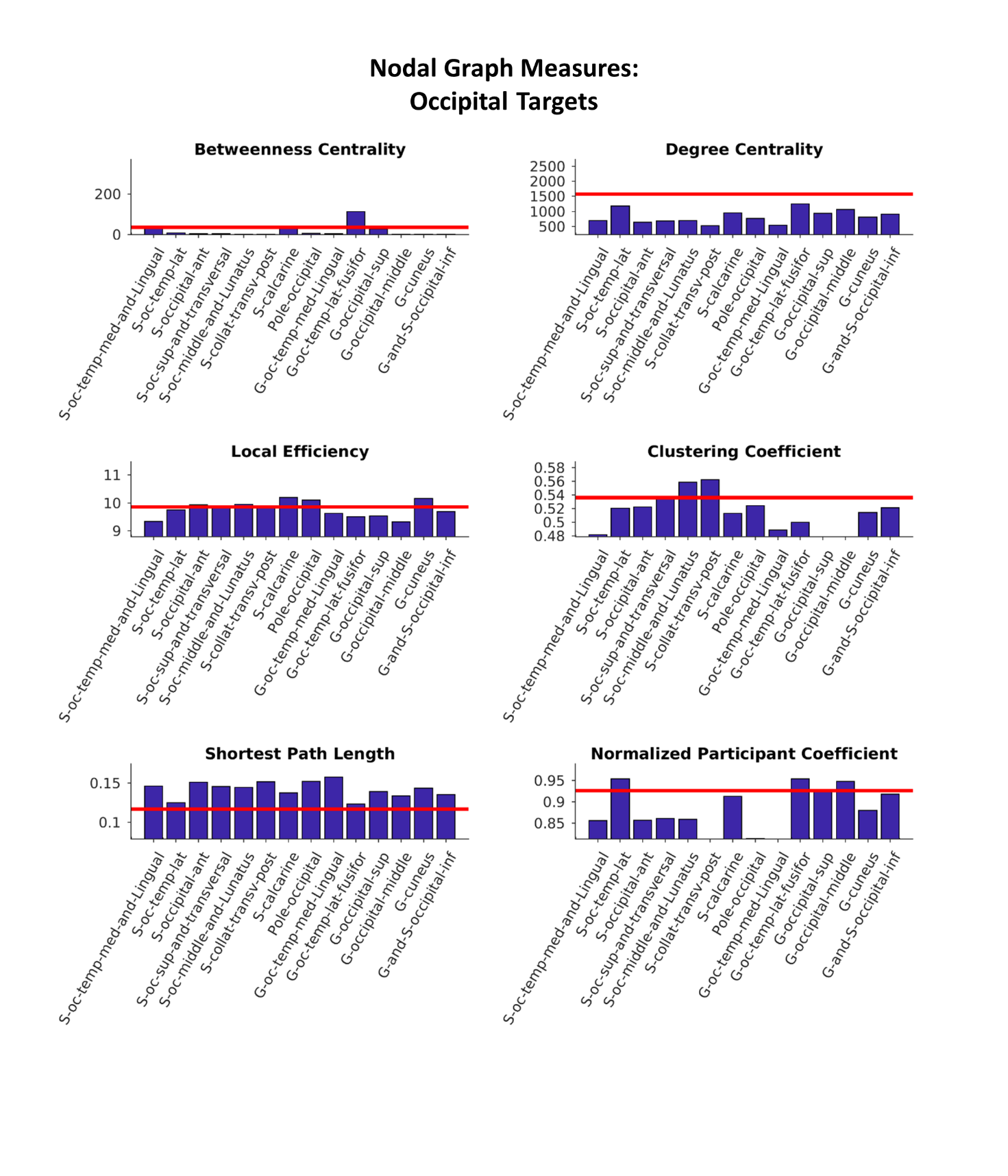

**Figure S11)** **Nodal graph-analysis metrics of the functional connectivity of targets of the occipital lobe**. Red lines mark the whole-network-averaged values.

### Autonomic-Vestibular Interactions

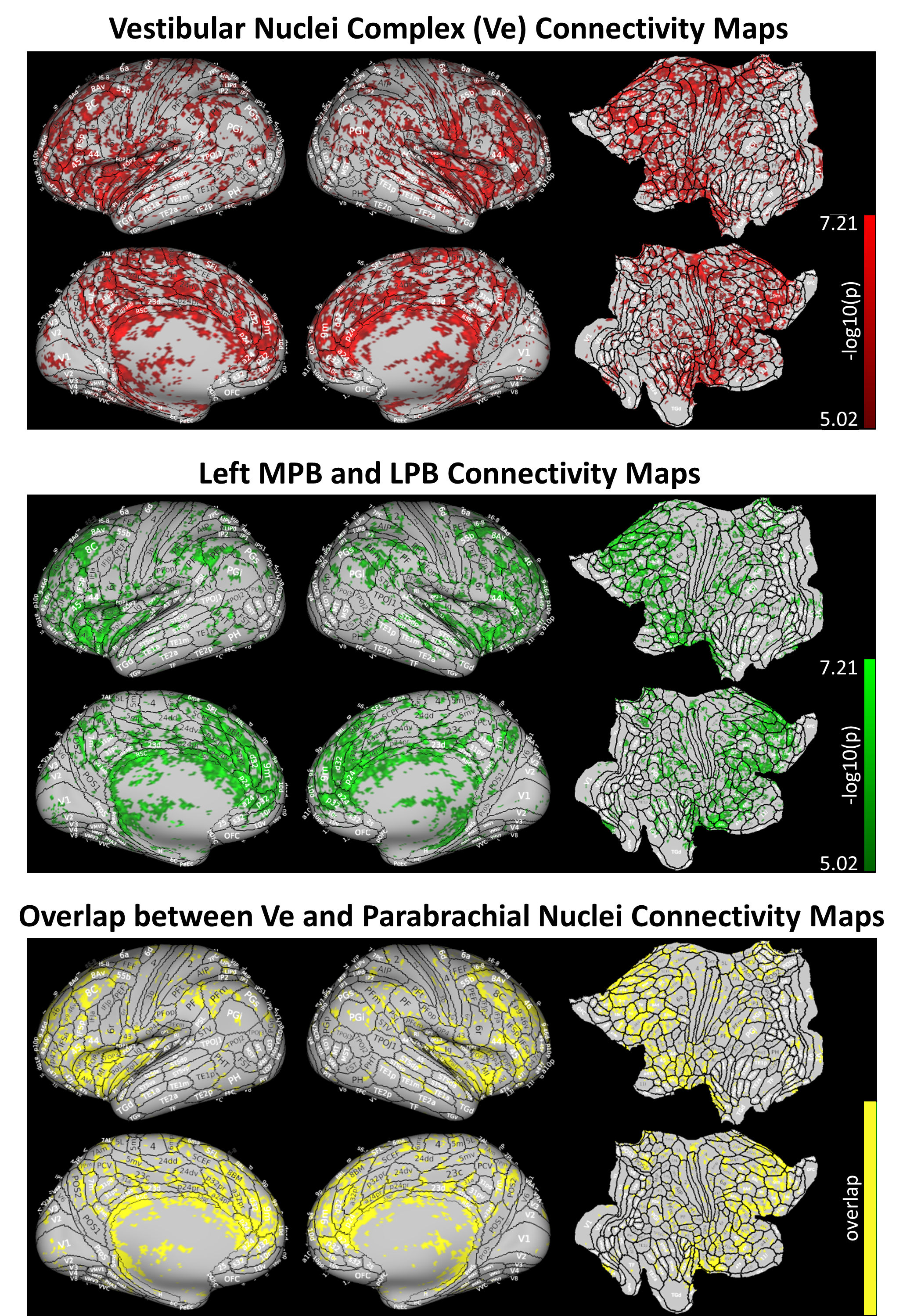

**Figure S12) Functional connectivity maps of autonomic and vestibular nuclei, and of their interaction, displayed on a flattened Glasser brain.** A) Voxel-based connectivity map (-log_10_(p-value), voxel-wise p<0.00001, cluster-wise p<0.01) obtained by group level regression analysis, using as seed timeseries the average signal extracted from the left Ve mask. B) Voxel-based connectivity map (-log_10_(p-value), voxel-wise p<0.00001, cluster-wise p<0.01) obtained by group level regression analysis, using as seed timeseries the average signal extracted from the left LPB and MPB masks: the union of the two connectivity maps is displayed. C) Regions of overlap between significant connectivity of left Ve, and the union of significant connectivity of left MPB and left LPB (voxel-wise p<0.00001, cluster-wise p<0.01). All maps are overlayed on the Glasser Parcellation [Glasser et al., 2016].
